# The latent stage of *Toxoplasma gondii* is targeted by the immune response and host protective

**DOI:** 10.1101/2024.03.05.583527

**Authors:** Lindsey A. Shallberg, Julia N. Eberhard, Aaron Winn, Sambamurthy Chandrasekaran, Christopher J. Giuliano, Emily F. Merritt, Elinor Willis, David A. Christian, Daniel L. Aldridge, Molly Bunkofske, Maxime Jacquet, Florence Dzierszinski, Eleni Katifori, Sebastian Lourido, Anita A. Koshy, Christopher A. Hunter

**Affiliations:** Department of Pathobiology, School of Veterinary Medicine and University of Pennsylvania; Philadelphia, PA 19104, USA; Department of Physics and Astronomy, School of Arts and Sciences, University of Pennsylvania; Philadelphia, PA 19104, USA; BIO5 Institute, University of Arizona; Tucson, AZ 85721, USA; Department of Immunology and University of Arizona; Tucson, AZ 85721, USA; Department of Neurology, University of Arizona; Tucson, AZ 85721, USA; Whitehead Institute for Biomedical Research; Cambridge, MA 02142, USA; Department of Biology, Massachusetts Institute of Technology; Cambridge, MA 02142, USA; Comparative Pathology Core, Department of Pathobiology, School of Veterinary Medicine, University of Pennsylvania; Philadelphia, PA 19104, USA; The Royal Ottawa Mental Health Center, Institute of Mental Health Research; Ottawa, Ontario, K1Z 7K4, Canada

**Author notes:** These authors contributed equally to this work (Lindsey A. Shallberg, Julia N. Eberhard, Aaron Winn).

## Abstract

Latency is a microbial strategy for persistence. For *Toxoplasma gondii* the ability of the bradyzoite stage to form long-lived cysts is critical for transmission, while their presence in neurons is considered important for immune evasion. Development of a mathematical model highlighted that immune pressure on bradyzoites should contribute to dynamics of cyst formation and reactivation. Experimental data demonstrated that a cyst-derived antigen was recognized by CD8^+^ T cells and that IFN-γ signaling in neurons contributes to cyst control. In addition, modeling and the use of a parasite strain unable to form bradyzoites revealed that this stage was not required for long-term persistence, but the absence of cyst formation resulted in increased tachyzoite replication in the CNS with associated tissue damage and mortality. Thus, the latent form of *T. gondii* is under immune pressure, mitigates infection-induced damage, and promotes survival of host and parasite.

## Introduction

Many infections can lead to an asymptomatic chronic state associated with low levels of micro-organisms despite the presence of protective immunity. For a subset of diverse viral and parasitic pathogens, persistence is linked to transition from a lytic to a non-lytic, latent developmental state. These quiescent stages are difficult to treat, and the absence of tractable models of natural host-pathogen interactions has left a gap in our understanding of the role of latency in immune evasion and persistence and its impacts on human health^1,2^. Latency is considered an evolutionarily conserved strategy for immune evasion that aids in persistence, while periodic reactivation is associated with disease and increased levels of transmission^3–8^. For example, the Sarcocystidae family of apicomplexan parasites (which includes *Toxoplasma gondii*) converts from the tachyzoite stage to the slow growing bradyzoite stage which forms long-lived tissue cysts found predominantly in neurons^6,9–11^. The molecular basis for this switch is mediated by the transcription factor BFD1 and the RNA binding protein BFD2 that enforce and maintain the bradyzoite transcriptional program^12–14^. This latent stage facilitates oral transmission and has been considered important for immune evasion, making it a major contributor to the evolutionary success of this parasite^15^.

*T. gondii* provides a system to study natural host-pathogen interactions in mice, and its genetic tractability has led to the development of a diverse toolkit to dissect how the immune system promotes long-term resistance to this organism^16,17^. The acute phase of infection is dominated by systemic dissemination of the lytic tachyzoite stage, which invades and replicates in many cell types and is associated with significant tissue damage. While this early phase of infection is controlled by the immune system, the chronic phase is characterized by conversion to the bradyzoite stage and the persistence of cysts predominantly within neurons of the central nervous system (CNS)^10,11^. While the cyst is largely considered dormant and the cyst wall impermeant, its relationship with the host cell is more dynamic than previously thought^18–20^. Cyst burden and size have been shown to change over time, and the ability of cysts to undergo periodic reactivation contributes to recrudescence and active disease in patients with acquired immune deficiencies that affect T cell function^11,21–23^. Resistance to *T. gondii* is marked by a balance between the ability to limit the tissue damage that results from tachyzoite replication versus the immune pathology associated with this infection. Thus, there are numerous host tolerance mechanisms that act to limit aberrant inflammation in the systemic phase of this infection as well as the immune pathology associated with the presence of *T. gondii* in the CNS^24–29^.

Many features of the brain contribute to its status as an immune privileged site^30,31^. Neurons have low basal levels of MHC class I and a reduced ability to respond to IFN-γ, features that have led to the concept that these long-lived cells provide a refuge for certain viral and parasitic pathogens^31–34^. This perspective is challenged by reports of CD8^+^ T cell recognition and killing of virally-infected neurons and evidence of non-cytopathic IFN-γ-mediated mechanisms of viral clearance from neurons^35–39^. Previous reports suggested that IFN-γ did not promote control of tachyzoites in neurons, however, more recent work has highlighted that extended incubation of neurons with IFN-γ limits parasite growth^34,40^. IFN-γ is considered the major mediator of resistance to *T. gondii*, but there is also evidence that a CD8^+^ T cell-mediated, perforin-dependent mechanism contributes to parasite control in the CNS and that CD8^+^ T cells can recognize neurons infected with tachyzoites^41–44^. Evidence that neurons containing cysts are directly targeted by CD8^+^ T cells is lacking, but there are reports that microglia and CD8^+^ T cells can interact with the cyst stage^42,45–49^. In mouse models, infection dynamics are characterized by a peak of cysts in the CNS in the first 3-5 weeks of infection and an increase in cyst size followed by a slow decline in cyst numbers^18,21,50,51^. Whether periodic cyst reactivation provides the immune response the opportunity to eliminate new tachyzoites and thereby limit formation of new cysts or there is a more direct immune-mediated mechanism to eliminate bradyzoites is unclear.

The ability to quantify T cell responses and changes in parasite burden during murine toxoplasmosis has been used as the basis for mathematical models to describe different facets of immunity to this infection^52–55^. Here, a series of ordinary differential equations (ODE) were developed to integrate how the immune response to different developmental stages of *T. gondii* might influence parasite dynamics. This model suggested the presence of immune-mediated cyst control and that cycles of cyst formation and reactivation are important for oscillations in parasite burden and CNS inflammation. To test these predictions, a combination of 1. transgenic parasites with stage-specific expression of the model antigen OVA, 2. parasites unable to form cysts, and 3. mice with blunted neuronal IFN-γ signaling were used. Together, these data reveal that an immune response is elicited against the latent form of *T. gondii,* and while this stage is not essential for persistence, the cyst provides a replicative sink required to mitigate infection-induced damage and thereby promote mutual survival of host and parasite.

## Results

### Modeling of parasite dynamics in the CNS predicts the presence of immune pressure on tachyzoites and bradyzoites

To investigate the relationship in the CNS between tachyzoite replication, cyst formation, reactivation, and the immune response, a system of ordinary differential equations (ODEs) was generated. A detailed derivation of the ODE system, explanation of the rate constants, and comparisons to the previous model are given in Supplementary Text section 1. The approach utilized is similar to that of Sullivan et al.^52^ but considers the possibility of immune pressure on bradyzoites. Two key assumptions of this model are that after tachyzoite entry to the CNS most of the host cells remain uninfected, and parasites spend the majority of their lifetime within infected cells. Important parameters are the numbers of tachyzoite-infected cells (*I_T_)*, bradyzoite-infected cells (*I_B_)*, and immune cells (Z). The number of tachyzoite-infected cells increases at a rate *β_T_* while the differentiation of tachyzoites to bradyzoites occurs at a rate *c_TB_*. The absence of cyst formation would yield a *c_TB_* = 0. While individual cyst growth *in vivo* is known to be asynchronous^18^, the parameter here considers the growth rate of the whole population as being synchronized. Bradyzoite reactivation is represented by the ability of bradyzoites to re-infect cells and give rise to tachyzoites at a rate *β_B_*. Cells infected with either tachyzoites or bradyzoites may rupture (at a rate *d_T_* or *d_B_*) or be cleared in the presence of immune cells (at a rate *ψ_T_ or ψ_B_*). Tachyzoite- or bradyzoite-infected cells trigger the immune response at a rate *a_T_* or *a_B_*. Here, the feedback between the immune response and infection takes the form of a predator-prey interaction: in the absence of an immune response parasite growth is unrestricted, but when immune cells are produced they limit parasite replication or lead to the elimination of infected cells. With decreased parasite burden, immune cells contract at a rate *µ*. Note that the nonlinear terms are normalized by the initial total uninfected cell number *S*_0_ (which essentially remains constant in our experiment) such that all rates are intensive. The system can be represented as a Petri net^56^ with compartments shown as circles and reactions as squares (Fig. 1a). In this visualization, the two arrows from *β_T_* to *I_T_* represent a +1 increase in parasite numbers and similarly these two arrows are also used to represent the ability of infected cells (*α_T_/α_B_*) to amplify the immune response (Z). The corresponding system of ODEs is given in equations (1), (2), and (3).

**Fig. 1:**
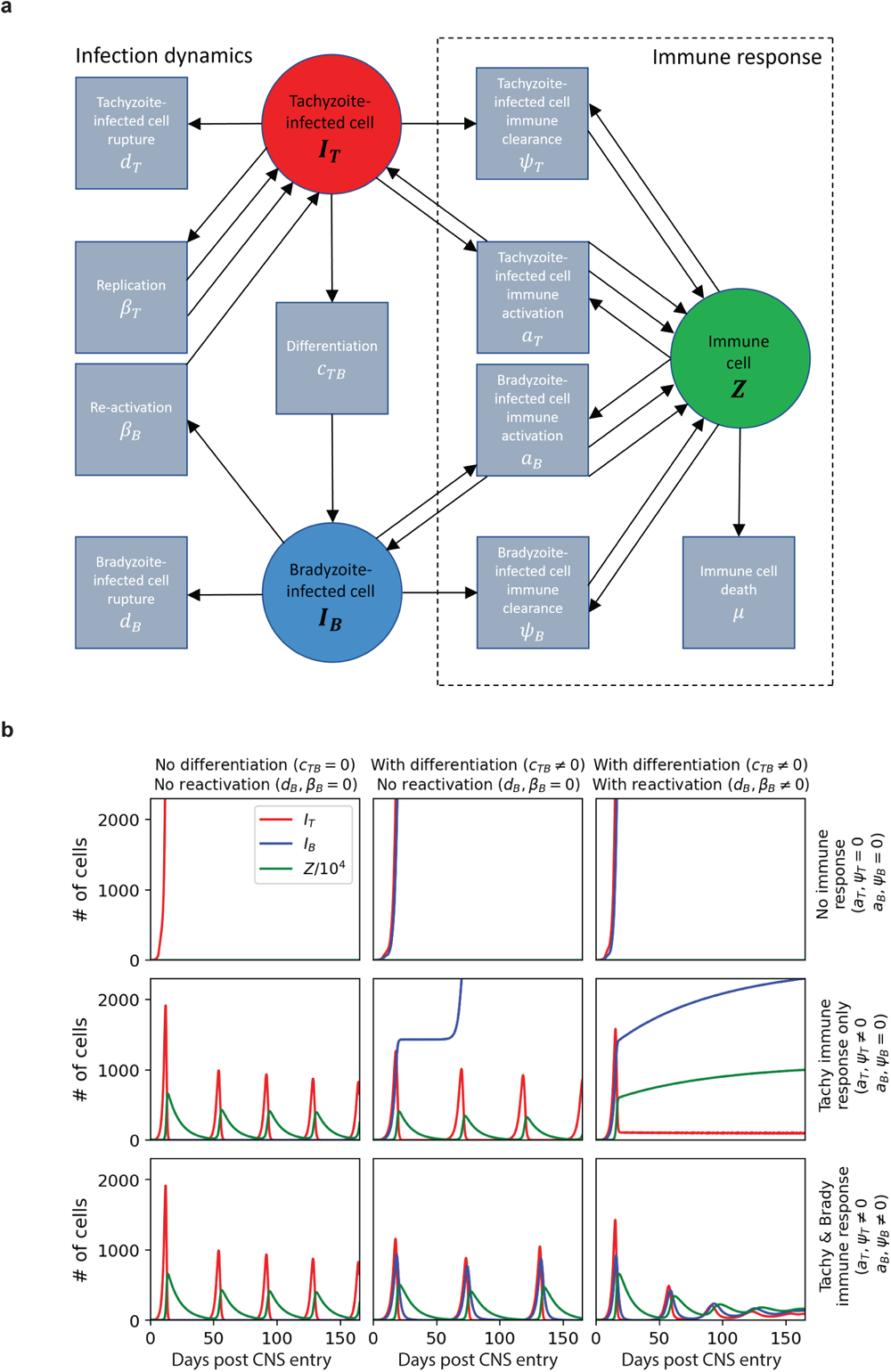
Compartmental modeling of *T. gondii* infection in the CNS predicts the presence of immune responses to tachyzoites and bradyzoites. **a**, Petri net representing the dynamics in equations (1), (2), and (3). Each square represents a particular reaction, with arrows entering a square representing reactants and arrows leaving a square representing products. **b**, Nine numerical results to (1), (2), and (3). Certain parameters are set to zero to demonstrate the role of each term. When nonzero, the parameters used were *S*_0_ = 10^8^, *I_T_* (0) = 1, *I_B_*(0) = 0, *Z*(0) = 10^5^, *β_T_* = 1.7, *β_B_* = .2, *c_TB_* = .25, *a_T_* = 10^5^, *a_B_* = .2 *∗* 10^5^, *µ* = .1, *ψ_T_* = 50, *ψ_B_* = 10.

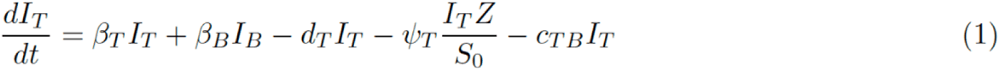

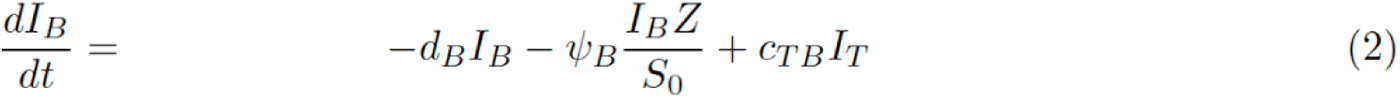

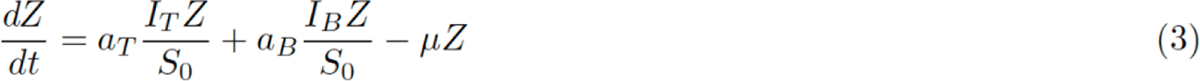

Primary data from murine infection were used to estimate model parameters (Supplementary Text section 4), and a summary is presented in Table 1. These biologically derived values were used to produce numerical solutions to equations (1), (2), and (3). By setting certain parameters equal to zero, the role of particular mechanisms can be elucidated, and several key scenarios are illustrated (Fig. 1b). In the top row, the immune response is disabled, and the number of tachyzoite-infected cells grows exponentially, and this remains largely unaltered even if bradyzoite differentiation and reactivation are incorporated. The solutions can be found analytically in this limit (see Supplementary Text section 2). In the second row, the inclusion of an anti-tachyzoite response alone (*ψ_T_*) results in the initial exponential growth of infected cells that triggers an influx of immune cells that leads to a rapid reduction in infected cells and contraction of the immune response. This is followed by cycles of tachyzoite recrudescence and control, apparent as oscillations in immune cell number. The inclusion of bradyzoite differentiation in the absence of reactivation (middle panel) predicts a larger period for the oscillations in tachyzoite-infected cell number but a stepped increase in bradyzoite numbers with each reactivation event. The inclusion of bradyzoite differentiation and reactivation in the absence of an anti-cyst response results in monotonic growth of the cyst burden (right hand panel). This simulation predicts the maintenance of immune populations which prevent significant oscillations in tachyzoite numbers. None of these scenarios in rows 1 or 2 capture all the key features of a natural infection in the CNS.

**Table 1:** Estimated parameter values based on empirical data in day^-1^ (Supplemental text).

| Parameter | Description | Value |
| --- | --- | --- |
| $d_T$ | Death rate of tachyzoite infected cells (without immune response) | 1 |
| $d_B$ | Death rate of bradyzoite infected cells (without immune response) | 0.01 |
| $\beta_T$ | Rate in which tachyzoites infect new cells | 1.7 |
| $\beta_B$ | Bradyzoite to tachyzoite reactivation rate | 0.2 |
| $c_{TB}$ | Tachyzoite to bradyzoite conversion rate | 0.25 |
| $a_T$ | Growth of tachyzoite induced immune pressure | $10^5$ |
| $a_B$ | Growth of bradyzoite induced immune pressure | $2 \cdot 10^4$ |
| $\mu$ | Contraction of immune response | 0.1 |
| $\psi_T$ | Immune pressure on tachyzoite infected cells | 50 |
| $\psi_B$ | Immune pressure on bradyzoite infected cells | 10 |
| $S_0$ | Initial cells susceptible to infection | $10^8$ |
| $I_T(0)$ | Initial number of tachyzoite infected cells | 1 |
| $I_B(0)$ | Initial number of bradyzoite infected cells | 0 |
| $C(0)$ | $I_T(0) + I_B(0)$ | 0 |
| $Z(0)$ | Initial number of immune cells | $10^5$ |

In the final set of simulations (bottom row), tachyzoite and bradyzoite specific immune responses are now incorporated – and in scenarios where differentiation to the bradyzoite is absent (left panel) this is followed by cycles of tachyzoite recrudescence and control that are similar in magnitude over the course of infection. However, when bradyzoite differentiation and reactivation are integrated (*β_B_* = 0 or *β_B_ ≠* 0) these oscillations are reduced in frequency and dampened over time (bottom right-hand panel). A stability analysis in Supplementary Text section 3 demonstrates that both differentiation and reactivation are necessary in order to dampen oscillations in infected cell number. It is shown that when *c_TB_* = 0 the oscillations in infected cell number are undampened, and when *c_TB_* is small but non-zero the oscillations decay at a rate proportional to *β_B_*. This model implies that both differentiation and reactivation are necessary to dampen oscillations in infected cell number and immune cell infiltration. While both rupture and immune clearance can lead to a decline in cyst numbers, rupture (and the ability to form new cysts) leads to a steady value of cysts, whereas immune clearance leads to cyclic changes and an overall decline in cyst populations. Together, these models predict that to recapitulate the key features of this infection in the CNS there needs to be immune mechanisms that exert pressure on both tachyzoite and bradyzoite stages.

### Cyst-derived antigen induces a CD8^+^ T cell response

To determine if cyst-derived antigen could be recognized by CD8^+^ T cells, transgenic parasites were generated with a cyst-specific promoter to drive expression of the model antigen OVA (bag1-OVA), and infection with bag1-OVA parasites was combined with intravenous (i.v.) transfer of T cell receptor (TCR) transgenic CD8^+^ OT-I T cells. The immune response to these parasites was compared to parasites that express OVA constitutively (by tachyzoites and bradyzoites) under the tubulin promoter (tub1-OVA; Fig. 2a). In C57BL/6 mice, during the acute phase the two parasite strains established similar levels of infection and induced comparable parasite-specific endogenous CD4^+^ and CD8^+^ T cell responses in the spleen (Extended Data Fig. 1a,b). As expected, only tub1-OVA parasites induced an endogenous SIINFEKL-specific response at this timepoint (Extended Data Fig. 1a). Infection was also performed one day post-transfer of congenically distinct CD45.1^+^CD45.2^+^ OT-I T cells that express Nur77^GFP^ upon recent TCR engagement (Extended Data Fig. 1c)^47^. OT-I T cells expanded and trafficked to the brain and spleen at 14 days post-infection (dpi) with tub1-OVA parasites, but this was not observed with bag1-OVA parasites. However, by 28 dpi, a timepoint when cyst formation is apparent in the CNS, both infections resulted in OT-I T cell populations in the brain (Extended Data Fig. 1e). These populations contained a subset of Nur77^GFP+^ cells, indicating that these T cells had received recent TCR stimulation (Extended Data Fig. 1f). The magnitude of the OT-I response in the CNS was approximately 5-to 10-fold greater during infection with tub1-OVA parasites compared to bag1-OVA parasites (Extended Data Fig. 1e). This difference could reflect delayed priming as a result of delayed OVA expression in bag1-OVA parasites. To normalize for time of priming, naive OT-I T cells were transferred into infected mice at 21 dpi (when both parasite strains are expected to express OVA) and analyzed 2-3 weeks later (Fig. 2b). The OT-I T cell response induced by the bag1-OVA parasites remained lower in magnitude and there was a lower frequency of Nur77^GFP^ expressing cells (Fig. 2c,d). These data suggest that the kinetics of T cell priming do not readily account for differences in the magnitude of the OT-I T cell response or the levels of TCR engagement observed between tub1-OVA and bag1-OVA parasites.

**Fig. 2:**
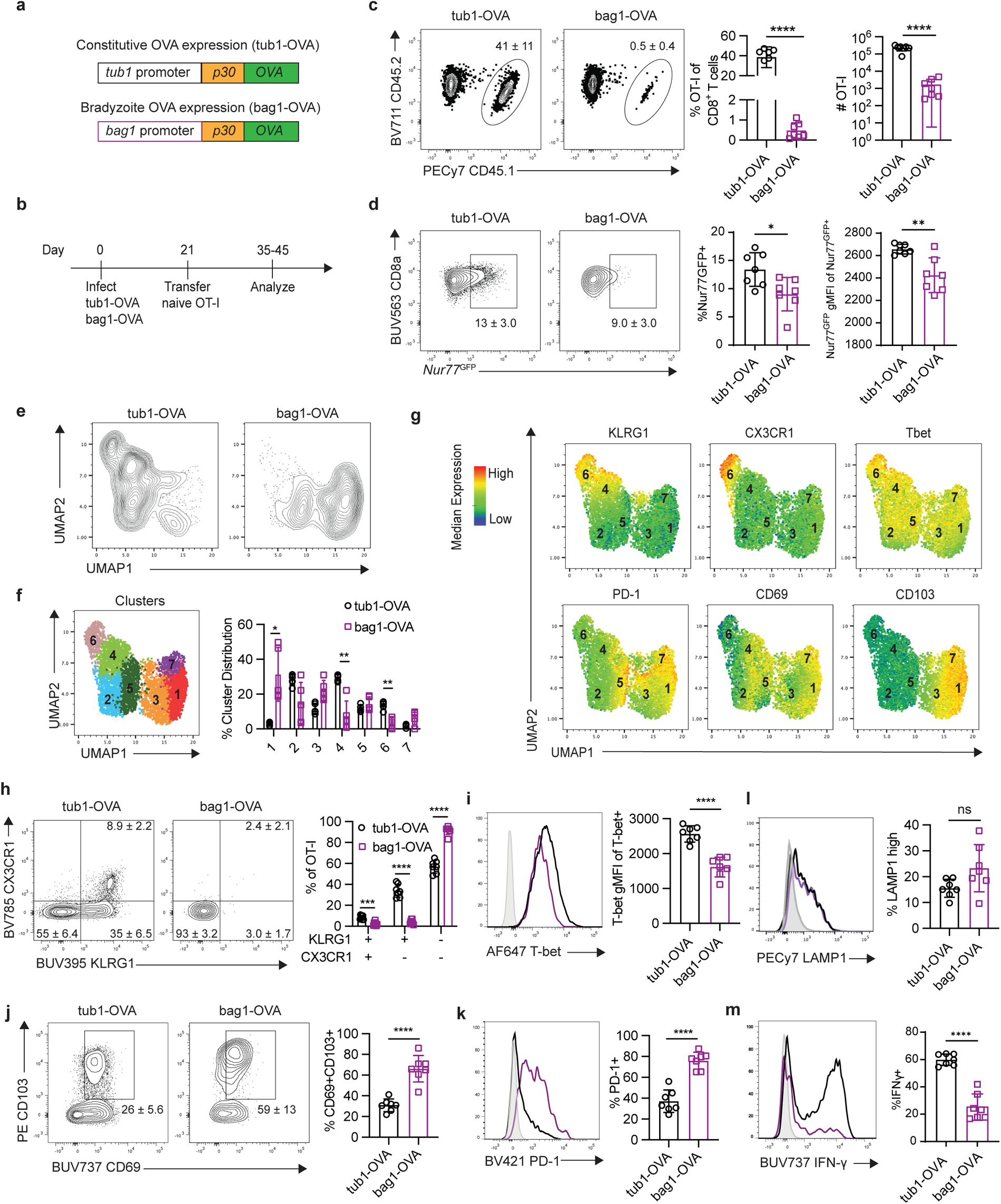
Cyst-derived antigen induces a CD8^+^ T cell response in the CNS. **a**, Transgenic parasite constructs for constitutive (tub1-OVA) and bradyzoite (bag1-OVA) restricted OVA expression. **b**, Experimental design for **c-m**. WT mice were infected intraperitoneally (i.p.) with either tub1-OVA (black, open circles) or bag1-OVA (purple, open squares) parasites. At 21 days post-infection (dpi) naïve congenically distinct (CD45.1^+^CD45.2^+^) Nur77^GFP^ OT-I T cells were transferred intravenously (i.v.). Brains were harvested and analyzed by flow cytometry 35-45 dpi. **c**, Frequency and number of OT-I T cells isolated from the brain of tub1-OVA or bag1-OVA infected mice. **d**, Frequency and geometric mean fluorescence intensity (gMFI) of Nur77^GFP^ expression in OT-I T cells shown in **c**. **e,f,** UMAP analysis and unsupervised clustering of OT-I T cells pooled from tub1-OVA and bag1-OVA infected brains. Colors represent 7 individual clusters identified through X-shift clustering analysis. **g**, Heatmaps displaying MFI of individual phenotypic markers across the 7 clusters identified in **f**. **h-k**, Flow cytometric profiling of OT-I T cells based on the frequency and gMFI of phenotypic marker expression. Grey histogram sample indicates expression by naive CD8^+^ T cells. **l, m**, OT-I degranulation and cytokine production measured by flow cytometry after 4 hour restimulation with SIINFEKL peptide. Grey indicates unstimulated OT-I T cell controls. Data are representative plots from 3 independent experiments with 3-7 mice per group. Bar graphs depict the mean ± SD. Data analyzed by two-tailed unpaired Student’s *t*-test. Bonferroni-Dunn correction for multiple comparisons included in statistical analyses performed in **f** and **h**. ns *p* > 0.05, \**p* < 0.05, \*\**p* < 0.01, \*\*\**p* < 0.001, and \*\*\*\**p* < 0.0001.

We next assessed the phenotype of OT-I T cells in the CNS responding to tub1-OVA or bag1-OVA parasites using high parameter flow cytometry. Uniform Manifold Approximation and Projection (UMAP) analysis of the concatenated OT-I T cell populations (from tub1-OVA and bag1-OVA infections) revealed 7 main clusters with varied levels of expression of effector T cell markers (KLRG1, CX3CR1, T-bet), inhibitory receptors (PD-1), and tissue residency markers (CD69, CD103) (Fig. 2e-g). OT-I T cells induced in response to tub1-OVA displayed heterogenous T effector phenotypes that contained KLRG1^+^CX3CR1^+^, KLRG1^+^CX3CR1^-^, and KLRG1^-^CX3CR1^-^ populations that expressed high levels of T-bet (Fig. 2h,i). Only a small proportion of cells were CD69^+^CD103^+^ or PD-1^hi^ (Fig. 2j,k). By contrast, OT-I T cells induced in response to bag1-OVA infection were dominated by a KLRG1^-^CX3CR1^-^ population with decreased T-bet expression. These cells also exhibited co-expression of tissue resident memory markers CD69 and CD103 and expressed high levels of the inhibitory receptor PD-1 (Fig. 2h-k). Restimulation of cells isolated from infected brains with the OVA-derived SIINFEKL peptide showed that while OT-I T cells from both infections had similar levels of degranulation (a measure of cytotoxic potential), T cells induced by bag1-OVA infection had a reduced ability to produce IFN-γ (Fig. 2l,m). A comparison of the OT-I T cell responses to tachyzoite-and bradyzoite-derived OVA indicates that tachyzoite-derived antigens drive the majority of the effector CD8^+^ T cell response in the CNS, but there is a sub-population of CD8^+^ T cells that respond to cyst-derived antigen, have a distinct phenotype, and reduced effector capacity.

### Neuronal expression of STAT1 is required for cyst control

The cytokine IFN-γ is the major mediator of resistance to *T. gondii* and signals through STAT1 to mediate its protective effects on haematopoietic and non-haematopoietic cell types ^44,57^. To test the role of IFN-γ in controlling *T. gondii* in neurons, *Snap25*-Cre mice were crossed with *Stat1^fl/fl^*mice to generate progeny (*Stat1^ΔNEU^*) in which neurons have a blunted ability to respond to IFN-γ. WT and *Stat1^ΔNEU^* mice were infected with tdTomato^+^ parasites to measure infected cells in the CNS. At 20 dpi, a timepoint prior to extensive cyst formation in the CNS, there was no difference between WT and *Stat1^ΔNEU^* mice in total parasite burden or the number of tdTomato^+^ leukocytes in the CNS (a proxy for tachyzoite-specific infection due to the absence of cysts in non-neuronal cells; Fig. 3a,b). At 3 months post-infection (mpi), total parasite burden remained comparable between WT and *Stat1^ΔNEU^* mice (Fig. 3c). However, at this timepoint, cysts in *Stat1^ΔNEU^*brains were larger (an indicator of increased longevity) and more numerous (Fig. 3d-f). This elevated cyst burden was not associated with increased mortality (Fig. 3g), and histopathological assessment of the brain did not reveal any signs of increased levels of CNS damage that would be associated with increased tachyzoite replication (Fig. 3h,i). In addition, there were no alterations in the numbers or phenotype of parasite-specific CD8^+^ T cells in the brains of these mice (Extended Data Fig. 2a-f). These results indicate that *in vivo* activation of neuronal STAT1 underlies a cell-intrinsic mechanism to control cysts. This observation is consistent with the modeling prediction that anti-cyst responses contribute to CNS parasite dynamics.

**Fig. 3:**
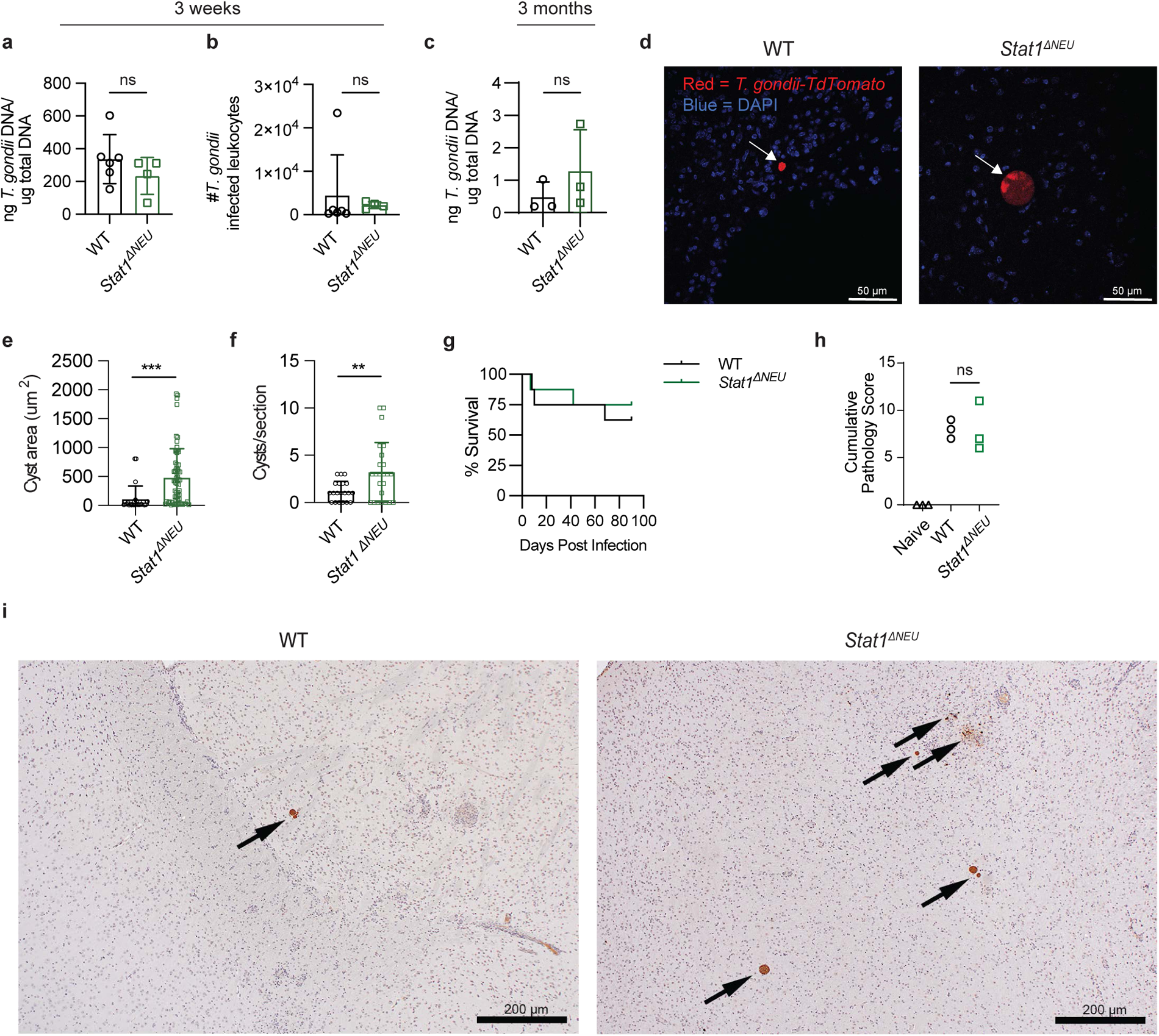
Neuronal STAT1 mediates cyst control during chronic *T. gondii* infection. WT and *Stat1^ΔNEU^* mice were infected i.p. with a tdTomato-expressing Pru strain of *T. gondii,* and brains were analyzed 3 weeks post-infection (wpi; **a,b**) or 3 months post-infection (mpi; **c-f,h,i**). **a,b**, Quantification of CNS parasite burden by qPCR (**a**) and tdTomato^+^ tachyzoite infected CD45^+^ leukocytes by flow cytometry (**b**) at 3 wpi. **c**, Quantification of CNS parasite burden at 3 mpi. **d**, Fluorescent imaging of brains from WT and *Stat1^ΔNEU^* mice at 3 mpi. (tdTomato^+^ parasites (Red), DAPI (blue), and white arrows indicate *T. gondii* cysts; scale bar = 50 μm). **e,f**, Brain cyst area (**e**) and number (**f**) at 3 mpi. Cysts were identified as vacuoles with >32 parasites present (5-20 cysts measured per section). **g**, Survival of infected WT and *Stat1^ΔNEU^* mice. **h**, H&E-stained sections of brain at 3 mpi were submitted for assessment and semiquantitative scoring by a board-certified veterinary pathologist. Cumulative pathological scores for individual mice are shown. **i**, IHC for *T. gondii* on sections from the brains scored in **h**. (Arrows indicate *T. gondii* cysts; scale bar = 200 μm). Data are representative of 2 independent experiments with 3-8 mice per group. Bar graphs depict the mean ± SD. Data analyzed by (**a-c,e,f**) two-tailed unpaired Student’s *t*-test or (**h**) Mann-Whitney test; ns *p* > 0.05, \**p* < 0.05, \*\**p* < 0.01, \*\*\**p* < 0.001.

### Cyst formation is not required for parasite persistence

We next modeled the impact of cyst formation and reactivation on parasite dynamics in the CNS. While inclusion of tachyzoite-to-bradyzoite conversion in the model produced infection dynamics that mirrored natural infection (Fig. 1), the inability to form bradyzoites (*c_TB_* = 0 versus *c_TB_* = 0.25) predicted an increase in the number of tachyzoite-infected cells early after parasite invasion of the CNS (Fig. 4a). In this scenario, parasite numbers would be controlled as infection progressed; however, even in the absence of cyst formation, low levels of tachyzoites would persist and undergo periodic recrudescence as immune cell numbers declined (Fig. 4b). To directly test this prediction, tub1-OVA parasites that lack BFD1 (Δ*bfd1*), a master regulator of tachyzoite to bradyzoite transition^12^, were generated (see Materials and Methods). At 14 dpi, mice infected with wild-type (WT) or Δ*bfd1* parasites had comparable parasite burden and T cell responses in the spleen as well as comparable systemic IFN-γ levels (Extended Data Fig. 3a-c). In contrast, higher levels of tachyzoite-infected cells were present in the brains of Δ*bfd1*-infected mice at 14 and 21 dpi. Interestingly, this was followed by a contraction from 30 to 45 dpi and a recrudescence at 60 dpi (Fig. 4c). Fluorescent imaging confirmed that the majority of parasitophorous vacuoles (PVs) in Δ*bfd1*-infected mice were negative for dolichos biflorus agglutinin (DBA) which binds the cyst wall, while all PVs in WT-infected mice were DBA^+^. (Fig. 4d,e). A fraction of Δ*bfd1* PVs were DBA^+^, but the DBA staining was weaker than what was seen with WT parasites, and the Δ*bfd1* parasites within DBA^low^ vacuoles were Sag1^+^ (a tachyzoite surface antigen) and SRS9^-^ (a bradyzoite surface antigen). Conversely, WT parasites within DBA^+^ vacuoles were Sag1^-^ and SRS9^+^ (Extended Data Fig. 3d). These data confirm that Δ*bfd1* parasites do not form cysts in the brain.

**Fig. 4:**
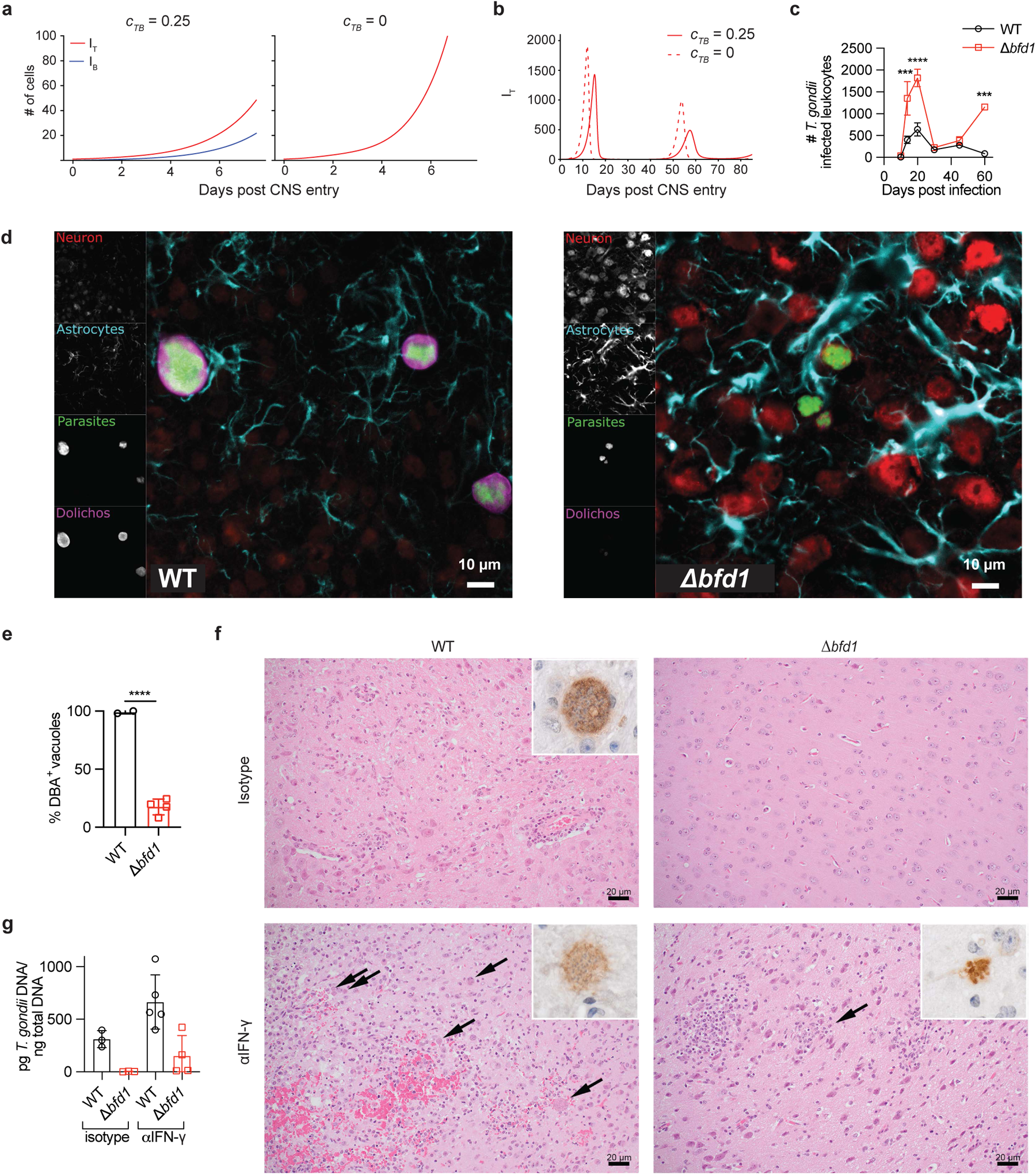
Latent stage conversion is not necessary for chronic infection. **a, b**, Model predicted CNS infection dynamics with or without parasite conversion from tachyzoites to bradyzoites (*c_TB_*). **a**, Early dynamics of CNS tachyzoite (I_T_, red) and bradyzoite (I_B_, blue) infected cell populations with cyst conversion (*c_TB_* = 0.25, left) or without (*c_TB_* = 0, right). **b**, Long-term dynamics of tachyzoite infected cell populations in the CNS with *c_TB_* = 0.25 (solid line) or *c_TB_* = 0 (dashed line). **c**, Quantification of tdTomato^+^ tachyzoite infected CD45^+^ leukocytes from the brains of WT (black, open circles) or Δ*bfd1* (red, open squares) infected mice, analyzed by flow cytometry. **d**, Fluorescent imaging of neurons, astrocytes, *T. gondii* parasites, and cysts in the brains of mice infected with WT (left) or Δ*bfd1* (right) parasites at 25 dpi. NeuN, MAP2, and Neurofilament (red); GFAP (cyan); tdTomato^+^ *T. gondii* (green); Dolichos biflorus agglutinin (DBA, magenta). Scale bar = 10 μm. n=2-4 mice per group, 19-200 vacuoles per mouse. **e**, Percentage normalization of DBA stained vacuoles at 25 dpi. **f, g**, Brains of mice infected with WT or Δ*bfd1* parasites were analyzed at 6 mpi, after treatment with 200 μg/dose of isotype or α-IFN-γ antibody 2 times/week for 4 weeks prior to harvest. **f**, Representative photomicrographs of brains from infected mice (H&E). Arrows indicate areas of *T. gondii* tachyzoite burden. Insets show cysts (upper left) or tachyzoites (lower left and right); IHC for *T. gondii.* Scale bar = 20 μm. **g**, Quantification of CNS parasite burden by qPCR. Data are representative of 1-2 independent experiments with 3-5 mice per group. Line graph depicts the mean ± SEM. Bar graphs depict the mean ± SD. Data analyzed by (**c**) 2-way ANOVA or (**e**) two-tailed unpaired Student’s *t*-test; \**p* < 0.05, \*\**p* < 0.01, \*\*\**p* < 0.001, and \*\*\*\**p* < 0.0001.

Since cyst formation is considered important for parasite persistence, it was possible that mice which survived infection with Δ*bfd1* parasites might fully clear the infection long-term. To determine if cyst formation is required for long-term parasite persistence, the brains of WT-and Δ*bfd1*-infected mice that had survived for 6 months were examined. Mice infected with WT parasites exhibited persistent encephalitis, apparent cysts, and detectable levels of parasite DNA by qPCR (Fig. 4f,g). In contrast, mice infected with Δ*bfd1* parasites lacked overt signs of ongoing inflammation or parasite replication, and parasite burden was below the level of detection by qPCR (Fig. 4f,g). However, because IFN-γ limits parasite replication in the CNS^58^, a cohort of mice at 6 mpi were treated with IFN-γ blockade for 4 weeks prior to harvest to determine if this would result in recrudescence. IFN-γ blockade resulted in a marked increase in parasite burden that was apparent by IHC and qPCR in the brains of both WT-and Δ*bfd1*-infected mice at 6 mpi. This increase was most prominent in WT-infected brains but was also observed in Δ*bfd1*-infected brains (Fig. 4f,g). Thus, while cyst formation helps maintain a higher parasite burden in the CNS, it is not essential for long-term persistence of *T. gondii*.

### Cyst formation promotes host protection from lethal tachyzoite replication

While mice infected with WT or Δ*bfd1* parasites had similar rates of survival between 10 and 14 dpi, those infected with Δ*bfd1* parasites showed increased mortality between 20 and 40 dpi (Fig. 5a). Treatment of Δ*bfd1*-infected mice with sulfadiazine (a drug that inhibits tachyzoite replication) starting at 21 dpi rescued mortality (Fig. 5a). Histopathological assessment of the brains of Δ*bfd1*-infected mice at 30 dpi showed areas of severe tissue necrosis surrounding foci of tachyzoite replication (Fig. 5b,c). In contrast, WT-infected brains, despite the presence of inflammation, glial reaction, and cysts, did not exhibit necrosis or remarkable damage to the tissue architecture (Fig. 5b,c). Differences in additional indicators of CNS pathology are also apparent in semiquantitative pathological grading of the brains of mice infected with WT and Δ*bfd1* parasites (Extended Data Fig. 5a,b). Together, these data indicate that higher mortality associated with Δ*bfd1* parasites is a consequence of increased levels of tachyzoite replication.

**Fig. 5:**
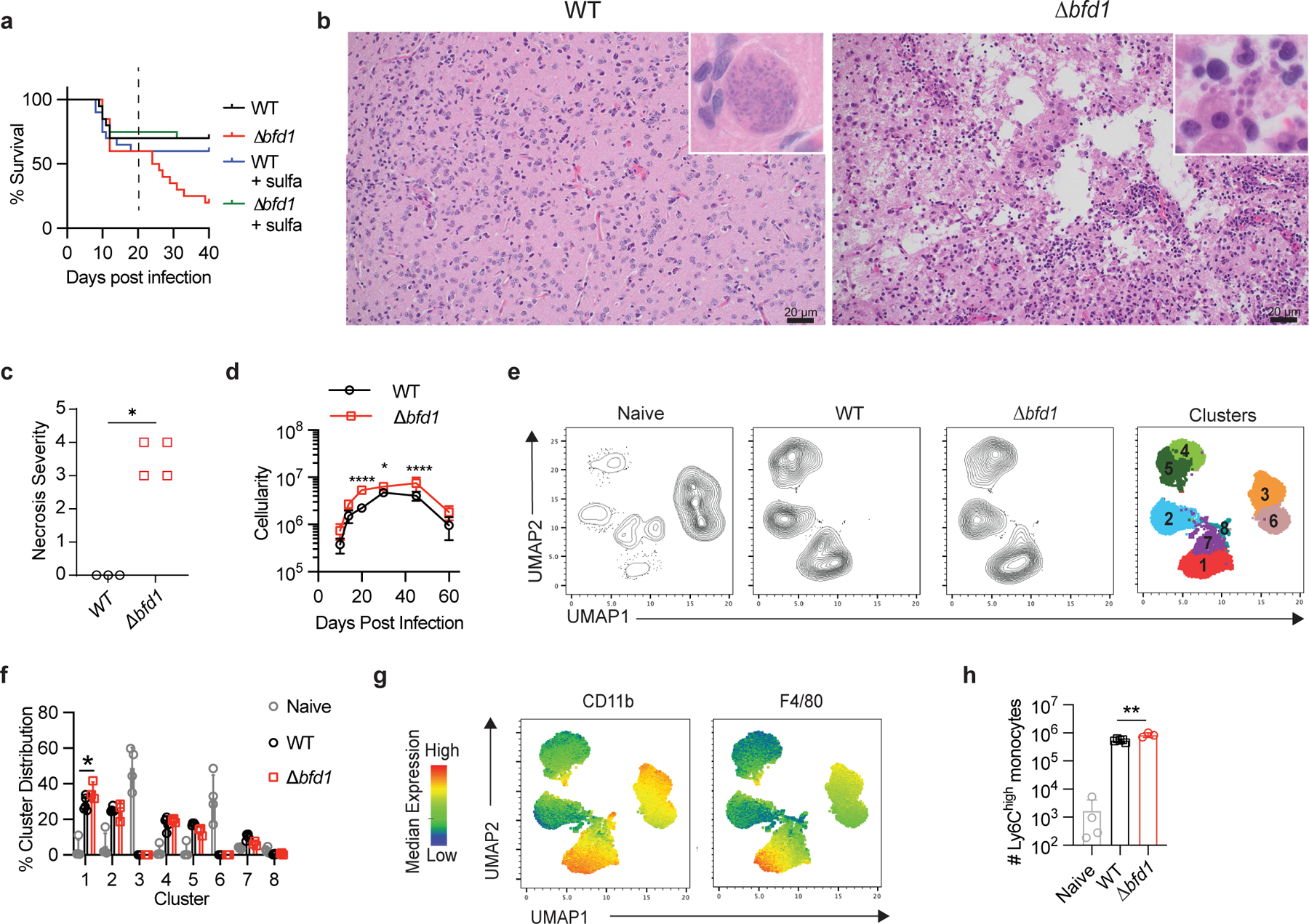
*Δbfd1* infected mice die of tachyzoite replication despite a competent immune response. **a**, Survival of mice infected i.p. with WT (black) or *Δbfd1* (red) parasites. A subset of mice received sulfadiazine treatment beginning at 21 dpi (WT=blue; *Δbfd1*=green). **b**, Representative photomicrographs of brains from mice infected with WT (left) or *Δbfd1* (right) parasites at 30 dpi. Insets show *T. gondii* cysts in WT infected brains and tachyzoites in *Δbfd1* infected brains. H&E; scale bar = 20 μm. **c**, H&E-stained sections of brain were submitted for assessment and semiquantitative scoring by a board-certified veterinary pathologist. Semiquantitative scores for necrosis severity shown. **d**, Brain leukocyte cellularity during chronic infection with WT (black, open circles) or *Δbfd1* (red, open squares) parasites was quantified by Guava ViaCount Assay. **e-h**, Flow cytometry analysis of the brains of naive mice (grey) and mice 30 dpi with WT (black) or *Δbfd1* (red) parasites. **e**, UMAP and unsupervised x-shift clustering analysis of CD45^+^ leukocytes from the brain at 30 dpi. **f**, Distribution of leukocytes amongst the 8 clusters shown in **e**. **g**, Heatmaps displaying MFI of phenotypic markers with the highest expression in cluster 1. **h,** Quantification of inflammatory monocytes in the brain (CD45^high^CD11b^+^F4/80^+^Ly6C^hi^). Data are representative of 2-3 independent experiments with 3-20 mice per group. Bar graphs depict the mean ± SD. Data analyzed by (**c**) Mann-Whitney test, (**d**) 2-way ANOVA, (**f**) two-tailed unpaired Student’s *t*-test with Bonferroni-Dunn correction for multiple comparisons, and (**h**) two-tailed unpaired Student’s *t*-test; \**p* < 0.05, \*\**p* < 0.01, \*\*\**p* < 0.001, and \*\*\*\**p* < 0.0001.

A possible explanation for increased tachyzoite replication during Δ*bfd1* infection could be a reduced immune response in the CNS. However, Δ*bfd1*-infected mice had increased numbers of immune cells in the CNS from 20 to 45 dpi compared to WT-infected mice (Fig. 5d, Extended Data Fig. 5c). By 60 dpi, CNS immune infiltration in Δ*bfd1*-infected mice had contracted to similar levels compared to WT-infected mice, a possible explanation for the resurgence in the number of Δ*bfd1* tachyzoite-infected cells shown previously at this timepoint (Fig. 5d, Fig. 4c). Phenotyping of the infiltrating immune cells revealed that the overall composition of the CNS immune response at 30 dpi was similar between WT and Δ*bfd1* infection (Fig. 5e,f, Extended Data Fig. 5d). The only apparent difference in the distribution of CNS immune cell populations was an increase in the proportion and number of inflammatory monocytes (CD45^hi^CD11b^+^F4/80^+^Ly-6C^hi^, cluster 1) during Δ*bfd1* infection (Fig. 5f-h). Phenotypic analysis was also performed in mice that received OT-I T cells one day prior to infection with WT or Δ*bfd1* parasites expressing tub1-OVA. At 14 and 30 dpi OT-I T cells in the brain were phenotypically similar between infections (Extended Data Fig. 5e), although at 30 dpi Δ*bfd1* infection was associated with a reduction in the CD69^+^CD103^+^ and KLRG1^-^CX3CR1^-^ memory-like populations (Extended Data Fig. 5f,g). Thus, the absence of cyst formation was not characterized by any overt loss of immune function. These results suggest that despite the presence of a competent immune response in the CNS, increased levels of tachyzoite replication in the brain during Δ*bfd1* infection leads to host mortality. Because the severity of toxoplasmic encephalitis (TE) varies with mouse strain, additional studies were performed in BALB/c mice, which have decreased neuropathology and enhanced repair compared to C57BL/6 mice^59,60^. Mortality was not observed in mice infected with either WT or Δ*bfd1* parasites (Extended Data Fig. 6a). However, BALB/c mice infected with Δ*bfd1* parasites showed enhanced immune infiltration, and the majority also showed increased pathology scores and the presence of necrosis in the CNS (Extended Data Fig. 6b-e). Additionally, the highest levels of parasite DNA were found in Δ*bfd1-*infected brains, similar to findings in C57BL/6 mice (Extended Data Fig. 6f). These data — in addition to a recent study showing higher tachyzoite burdens in CBA mice infected with a type III strain (VEG) of *T. gondii* lacking BFD2^14^ — highlight that across mouse models of varying resistance to TE, the inability of *T. gondii* to form cysts is associated with increased levels of tachyzoite replication and associated pathology in the CNS.

## Discussion

While neurons represent a cellular refuge from the immune system for many pathogens, there is evidence that neurons infected with viruses or *T. gondii* tachyzoites can be recognized by CD8^+^ T cells^35,43,61^. However, the role of CD8^+^ T cells in control of the latent stage of *T. gondii* is less well understood. There are reports that depletion of CD8^+^ T cells in chronically infected mice does not impact cyst numbers, whereas other studies have shown that CD8^+^ T cells contribute to reduced cyst burden. Perforin-deficient mice have increased cyst numbers, and the transfer of CD8^+^ T cells from infected mice led to reduced cyst numbers in infected SCID mice in a perforin-dependent manner^41,42,49,62^. In the latter studies, protective CD8^+^ T cell responses are directed against GRA6, a parasite antigen that is expressed at highest levels in tachyzoites, associated with the intravacuolar membrane, and activates the host transcription factor NFAT^63,64^. Because this antigen is expressed in both tachyzoites and bradyzoites, and there are reports that CD8^+^ T cell recognition of a tachyzoite-expressed model antigen can lead to decreased cyst numbers^43^, it is unclear whether the CD8^+^ T cell-mediated reduction in cysts is due to control of tachyzoites or direct lysis of neurons that contain cysts. Intravital imaging of the CNS has revealed CD8^+^ T cells arrested in the vicinity of cysts, but direct interactions with this structure and T cell–mediated lysis of bradyzoite-infected neurons have not been observed^45–47,54^. Altogether these studies implicate a role for CD8^+^ T cells in the reduction of *T. gondii* cysts, but direct evidence of immune responses targeting neuronal cysts has not been shown. Here, we demonstrate through (1) generation of CD8^+^ T cell responses against cyst-derived antigen and (2) increased cyst burden with ablation of neuronal STAT1 that there is immune-mediated recognition and control of *T. gondii* cysts. These findings, combined with the models of host-parasite dynamics in the CNS, suggest that a combination of tachyzoite clearance and bradyzoite-directed immune pressure is required to produce the progressive reduction in cyst numbers that is typical of this infection^21,50^.

These observations raise new questions about the mechanisms that underlie effector CD8^+^ T cell recognition of cyst-derived antigens. It is possible that the ability of neurons to upregulate MHC class I allows infected neurons to directly present cyst-derived antigen to CD8^+^ T cells. Alternatively, periodic cyst reactivation could release cyst-derived antigens and provide the opportunity for cross-presentation by microglia or dendritic cells present in the CNS during TE. Regardless of the mechanism of antigen presentation, we favor a model analogous to some viral systems where local production of IFN-γ provides sustained signals to help neurons control infection^36,38,39,65^. This mechanism of non-cytopathic IFN-γ mediated control may help explain the presence of neurons that had previously been infected with *T. gondii* but cleared the parasite^66^. Many distinct IFN-γ–mediated anti-microbial mechanisms have been shown to contribute to the control of tachyzoites across cell types and species^67^. However, though immunity-related GTPases have been implicated in the ability of a subset of neurons to clear *T. gondii,* the relevance of these same mechanisms to bradyzoite control is uncertain^34^.

While the focus of this study has been on the mechanisms involved in cyst control, the fact that cysts persist in the presence of immune pressure suggests that this latent stage can either evade IFN-γ–mediated anti-microbial activities or mitigate immune effector responses. Consistent with the former idea are reports that bradyzoites also produce the parasite virulence factor TgIST, which is known to antagonize STAT1 activity upon secretion by tachyzoites^68–70^. Expression of this STAT1 antagonist in bradyzoites would suggest that this stage is under selective pressure from IFN-γ. In addition, the finding that the CD8^+^ T cell response to cyst-derived antigen from bag1-OVA parasites comprises a phenotypically distinct sub-population of CD69^+^CD103^+^PD1^hi^CD8^+^ T cells was unanticipated. Previous reports have highlighted the presence of a T_rm_-like population of CD69^+^CD103^+^CD8^+^ T cells and expression of PD-1 on CD8^+^ T cells during TE^45,71,72^. PD-1 is an inhibitory receptor that limits T cell activation and has been linked to the idea that CD8^+^ T cells during chronic toxoplasmosis have reduced effector functions^73^. Thus, the PD-1^hi^ phenotype of the CD8^+^ T cells responding to bag1-OVA could underlie their reduced ability to produce IFN-γ. Because neurons express low levels of MHC class I, lack costimulatory molecules, and express PD-L1^74,75^; it is possible that class I–restricted CD8^+^ T cell interactions with infected neurons lead to sub-optimal T cell responses. Decreased effector capacity of T cells responding to infected neurons could underlie the attrition, but incomplete elimination, of cysts.

The use of ODE models to understand host-pathogen dynamics provides an opportunity to manipulate aspects of parasite biology and the host immune response in ways that are not always accessible through experimentation^76,77^. Others have noted the lack of tractable models makes it hard to test how latency impacts other pathogens^1^, but the statistical approaches described here can be adapted to incorporate strain or pathogen-specific features associated with *T. gondii* or other infections. For example, differences in the rate of cyst formation or ability of different stages to evade the immune response can be accounted for by adjusting the model parameters (*I_B_)* or (*ψ_T_* or *ψ_B_*), respectively. This combination of modeling and *in vivo* experiments provides new insight into the interplay between the latent stage of *T. gondii* and the host immune response. In particular, the data presented highlight that cyst development provides a replicative sink that tempers tachyzoite expansion in the CNS, mitigating damage to the host. This illustrates the concept that avirulence can be a key feature of the parasitic lifestyle, supported by the existence of defined developmental stages associated with mitigation of virulence across multiple parasites, including *Trypanosoma brucei* and Schistosomes^78–80^. Thus, *T. gondii* cyst formation in neurons represents a tradeoff between quiescence, which limits parasite replication, and delayed differentiation, which risks damage to the host.

## Materials and Methods

### Mice

Mice were housed in the University of Pennsylvania vivarium according to institutional guidelines. C57BL/6, CD45.1, *Nur77*^GFP^, OT-I, *Stat1*^Flox^, and BALB/c mice were purchased from Jackson Laboratories and bred at the University of Pennsylvania. *Snap25*^Cre^ mice were generously provided by Hongkui Zeng at the Allen Institute for Brain Science. All mice are on a C57BL/6 background unless otherwise noted. Ethical oversight of all animal studies was approved by the University of Pennsylvania Institutional Care and Use Committee.

### Infections and T cell transfers

*T. gondii* parasites were maintained in culture in human foreskin fibroblasts (HFF). To isolate tachyzoites for infection, *T. gondii*-infected HFFs were gently scraped from flasks and passed 5 times through a 26G syringe and washed with PBS. Mice were infected by i.p. injection with 5,000-10,000 tachyzoite parasites grown *in vitro*, unless otherwise stated.

For OT-I T cell transfers, whole splenocytes from Naive Nur77^GFP^ CD45.1^+^CD45.2^+^ OT-I mice were processed as described below. The fraction of OT-I cells in the splenocytes was determined by flow cytometry, and 5,000 OT-I cells were transferred i.v. into mice at the indicated timepoint prior to or post-infection.

### IFN-γ blockade and sulfadiazine treatment

*In vivo* blockade of IFN-γ signaling was performed by i.p. injection of 200 μg per dose of rat IgG1 anti-IFN-γ (clone XMG1.2, BioXcell) or IgG1 anti-horseradish peroxidase for control mice (HRPN, BioXcell). Injections were administered 2 times per week for 4 weeks prior to harvest and analyses for parasite burden.

Where indicated, mice were treated with the anti-parasitic drug Sulfadiazine (Sigma-Aldrich: S8626-25G) via drinking water. Sulfadiazine was reconstituted in dimethylsulfoxide (DMSO) to 50 mg/mL and added to drinking water at a final concentration of 0.25 mg/mL. Sulfadiazine treated water was administered starting at 21 dpi and refreshed every three days for 4 weeks.

### Tissue processing and cell counting

To generate single cell suspensions for flow cytometry, spleens were passed through a 40μm filter and red blood cells were lysed for 3 minutes at room temperature in ACK lysis buffer. Brains were diced into 1mm pieces and digested at 37C 5% CO_2_ for 1.5 hours with 250ug/mL collagenase/dispase and 10μg/mL DNase, and then passed through a 70μm filter. Leukocytes were then isolated through a 30% and 60% percoll gradient and density centrifugation at 2000rpm for 25 minutes. Whole blood was collected through submandibular bleed into 0.05mM EDTA in PBS. Cells were pelleted and red blood cells were lysed for 3 minutes at room temperature in ACK lysis buffer.

For quantification of cellularity and live leukocytes isolated from tissues, a fraction of processed cell suspensions were stained with Guava ViaCount Reagent (Cat. No. 11-25209, 240 mL) and analyzed on a Guava easyCyte Flow cytometer according to manufacturer protocol.

### Generation of transgenic parasites

To generate Δ*bfd1* parasites, Pru-tub1-OVA-tdTomato parasites were mechanically lysed from host cells by scraping and syringe releasing through a 27G needle. Parasites were pelleted for 10 minutes at 1000g, resuspended in Cytomix (10 mM KPO_4_, 120 mM KCl, 150 mM CaCl_2_, 5 mM MgCl_2_, 25 mM HEPES, 2 mM EDTA), and combined with a DNA transfection mixture to a final volume of 400μl. The final transfection mixture was supplemented with 2 mM ATP and 5 mM glutathione. Two gRNAs contained on a Cas9 expressing plasmid were transfected to target the regions immediately upstream and downstream of the BFD1 locus as previously described ^20^. A pTUB-mNeonGreen repair template with homology arms matching the 40 bp regions flanking the cut sites was transfected to allow for sorting of Δ*bfd1* parasites by FACS. After sorting, parasites were plated at limiting dilutions to allow for screening of clonal parasites. A Δ*bfd1* clone was confirmed by PCR and sanger sequencing. However, despite these parasites being mNeonGreen-positive, the mNeonGreen repair template was not found within the BFD1 locus, indicating that this construct had randomly integrated into the parasite genome.

Upstream gRNA sequence: GTTGAGTCCAAGCAGAGCTC

Downstream gRNA sequence: GTGTAGAGTCGTGGAAGGAG

BFD1 PCR primer forward (to confirm knockout): cctcatccttcgtcacgcgt

BFD1 PCR primer reverse (to confirm knockout): tgcttcgggcaggcgactat

Generation of *bag1*-OVA parasites was performed as described previously^11^: Vectors were generated to contain (i) the *bag1* promoter upstream of (ii) the last 31 amino acids of from the *T. gondii* major glycosyl-phosphatidylinositol-anchored surface antigen, P30, containing the signal sequence for targeting this protein to the parasitophorous vacuole, followed by (iii) amino acids 140-386 of *ovalbumin* and (iv) a 3’ untranslated region of the *dihydrofolate reductase-thymidylate synthase* gene. Transfections of type II Pruginaud (Pru) strain parasites were performed using electroporation, and stably expressing parasites were selected with chloramphenicol. Clonal parasite lines were generated by limiting dilution.

### Immunohistochemistry

Brains were dissected by a sagittal cut along the midline, collected in 10% formalin, embedded in paraffin, and sectioned. For identification of *T. gondii* parasites, slides were hydrated and antigen was retrieved with 0.01M sodium citrate buffer at pH 6.0, endogenous peroxidase was blocked with 0.3% H_2_O_2_, and sections were blocked with 2% goat serum. Parasites were detected with rabbit anti-*T. gondii* polyclonal antibody (gifted from Fausto Araujo, Palo Alto Medical Foundation, 1:1000) followed by biotinylated goat anti-rabbit IgG antibody (Vector Laboratories). ABC reagent and DAB substrate (Vector Laboratories) were used according to manufacturer’s protocol to visualize parasite staining, and hematoxylin staining was applied to visualize nuclei. Images were acquired with a Leica DM6000 Widefield Fluorescence Microscope.

### Pathological Assessment

Brains were collected and processed for sectioning as described above. Hematoxylin and eosin-stained sagittal sections of the brain, including the cerebral cortex and basal nuclei (Fig. 5 and Extended Data Fig. 5) or cerebral cortex and basal nuclei, hippocampus, thalamus, midbrain, cerebellum, pons, and medulla (all other assessments) were assessed by a board-certified veterinary pathologist. The type of inflammatory cells and semiquantitative scores for the severity of parameters of interest (inflammation, hemorrhage, gliosis, necrosis, and presence of parasites) were recorded for individual animals.

### Flow cytometry

Single cell suspensions were plated at up to 2x10^6^ cells. Cells were Fc receptor blocked for 20 minutes with 0.5μg/mL αCD16/32 (Clone 2.4G2) and 0.25% normal rat serum in FACs buffer (2% BSA and 0.02mM EDTA in PBS) at 4°C. If staining for tetramer, cells were then washed and incubated with 1:200-1:400 dilution of tetramer in FACs buffer at room temperature for 30 minutes. Following Fc blocking or tetramer stain, surface stain was applied and incubated for 30 minutes at 4°C. Cells were then washed and resuspended in 0.1% paraformaldehyde in FACs buffer. If staining for intracellular antigens, cells were fixed and permeabilized with eBioscience Foxp3 / Transcription Factor Staining Buffer Set according to manufacturer’s protocol. Cells were then stained for intracellular antigens in permeabilization buffer for 30 minutes at 4°C. Cells were washed and resuspended in FACs buffer. Stained cells were acquired on a BD LSR Fortessa, BD FACSymphony A5, or BD FACSymphony A3. Analysis was performed with FlowJo v10.7.2.

The following antibodies and reagents were used for staining: B220: BUV496, BD Biosciences: 612950, clone: RA3-6B2, RRID:AB_2870227; CD3: APC-ef780, Invitrogen: 47-0032-82, clone: 17A2, RRID:AB_1272181; CD3: BUV737, BD Biosciences: 612803, clone: 17A2, RRID:AB_2738781; CD3e: PE-cf594, BD Biosciences: 562286, clone: 145-2C11, RRID:AB_11153307; CD4: BUV496, BD Biosciences: 612952, clone: GK1.5, RRID:AB_2813886; CD4: BV650, Biolegend: 100555, clone: RM4-5, RRID:AB_2562529; CD4: FITC, eBioscience: 11-0041-85, clone: GK1.5, RRID:AB_464892; CD4: APC-ef780, Invitrogen: 47-0041-82, clone: GK1.5, RRID:AB_11218896; CD8a: BUV563, BD Biosciences: 748535, clone: 53-6.7, RRID:AB_2872946; CD8a: BUV615, BD Biosciences: 613004, clone: 53-6.7, RRID:AB_2870272; CD8a: BV650, Biolegend: 100742, clone: 53-6.7, RRID:AB_2563056; CD8b: APC-ef780, Invitrogen: 47-0083-82, clone: eBioH35-17.2, RRID:AB_2573943; CD11a: BUV805, BD Biosciences: 741919, clone: 2D7, RRID:AB_2871232; CD11b: BV650, Biolegend: 101259, clone: M1/70, RRID:AB_2566568; CD19: BUV395, BD Biosciences: 563557, clone: 1D3, RRID:AB_2722495; CD45: AF647, Biolegend: 103124, clone: 30-F11, RRID:AB_493533; CD45.1: ef450, Invitrogen: 48-0453-82, clone: A20, RRID:AB_1272189; CD45.1: BV711, Biolegend: 110739, clone: A20, RRID:AB_2562605; CD45.1: PE-Cy7, Biolegend: 110730, clone: A20, RRID:AB_1134168; CD45.2: APC, Biolegend: 109814, clone: 104, RRID:AB_389211; CD45.2: BV711, Biolegend: 109847, clone: 104, RRID:AB_2616859; CD69: BUV737, BD Biosciences: 612793, clone: H1.2F3; CD69: PerCP-Cy5.5, eBioscience: 45-0691-82, clone: H1.2F3, RRID:AB_1210703; CD103: PE, eBioscience: 12-1031-81, clone: 2E7, RRID:AB_11150242; CD103: BV605, Biolegend: 121433, clone: 2E7, RRID:AB_2629724; CD107a: PE-Cy7, Biolegend: 121620, clone: 1D4B, RRID:AB_2562146; CD127: BV421, Biolegend: 135027, clone: A7R34, RRID:AB_2563103; CTLA-4: APC-R700, BD Biosciences: 565778, clone: UC10-4F10-11, RRID:AB_2739350; CX3CR1: BV785, Biolegend: 149029, clone: SA011F11, RRID:AB_2565938; CX3CR1: PerCP-Cy5.5, Biolegend: 149009, clone: SA011F11, RRID:AB_2564493; F4/80: APC-ef780, eBiosciences: 47-4801-82, clone: BM8, RRID:AB_2735036; GFP: AF488, Biolegend: 338008, clone: FM264G, RRID:AB_2563288; H-2Kb: AF647, Biolegend: 116512, clone: AF6-88.5, RRID:AB_492917; I-A/I-E: AF700, Biolegend: 107622, clone: M5/144.15.2, RRID:AB_493727; I-A/I-E: BV711, Biolegend: 107643, clone: M5/144.15.2, RRID:AB_2565976; IFN-γ: BUV737, BD Biosciences: 612769, clone: XMG1.2; Ki-67: BV470, BD Biosciences: 566109, clone: B56, RRID:AB_2739511; KLRG1: BUV395, BD Biosciences: 740279, clone: 2F1, RRID:AB_2740018; Ly-6C: BV785, Biolegend: 128041, clone: HK1.4, RRID:AB_2565852; Ly-6G: BUV563, BD Biosciences: 612921, clone: 1A8, RRID:AB_2870206; NK1.1: BUV395, BD Biosciences: 564144, clone: PK136, RRID:AB_2738618; PD-1: BV421, Biolegend: 135221, clone: 29F.1A12, RRID:AB_2562568; PD-1: BV605, Biolegend: 135220, clone: 29F.1A12, RRID:AB_2562616; PD-1: BV785, Biolegend: 135225, clone: 29F.1A12, RRID:AB_2563680; T-bet: AF647, Biolegend: 644804, clone: 4B10, RRID:AB_1595466; TCRβ: PerCP-Cy5.5, Biolegend: 109228, clone: H57-597, RRID:AB_1575173; TCRβ: ef450, Invitrogen: 48-5961-82, clone: H57-597, RRID:AB_11039532; Tetramer MHCI (OVA): PE, NIH Tetramer Core, peptide: SIINFEKL; Tetramer MHCI (Tgd057): APC, NIH Tetramer Core, peptide: SVLAFRRL; Tetramer MHCII (AS15): APC, NIH Tetramer Core, peptide: AVEIHRPVPGTAPPS; Tim3: BV605, Biolegend: 119721, clone: RMT3-23, RRID:AB_2616907; TNF-α: APC, Invitrogen: 17-7321-82, clone: MP6-XT22, RRID:AB_469508; Vα2 TCR: BUV615, BD Biosciences: 751416, clone: B20.1, RRID:AB_2875415; Vα2 TCR: PE, Biolegend: 127808, clone: B20.1, RRID:AB_1134183; Viability: GhostDye Violet 510, TONBO Biosciences: 13-0870-T100; Viability: GhostDye Red 780, TONBO Biosciences: 13-0865-T100.

### T cell peptide restimulation

Whole splenocytes or brain leukocytes were plated at a constant cell concentration. Cells were incubated with 1μM SIINFEKL (OVA peptide), SVLAFRRL (Tgd057 peptide), or AVEIHRPVPGTAPPS (AS15 peptide) and fluorescently labeled αLAMP1 antibody for 2 hours, followed by a further 2 hours with Protein Transport Inhibitor Cocktail (Invitrogen: 00-4980-03). Cells were analyzed for degranulation and cytokine production by flow cytometry.

### Fluorescent imaging

Brains were harvested from mice following perfusion with heparin in saline followed by 4% paraformaldehyde and stored in 30% sucrose in saline, as described previously ^29^. Staining and imaging for neurons, astrocytes, and dolichos biflorus lectin for WT- and Δ*bfd1-* infected brains was performed as described previously^29^. The following reagents and antibodies were used: DBA (Vector laboratories B1035); GFAP (DAKO Z0334); MAP2 (Abcam ab5392); NeuN (Millipore MAB3778); Neurofilament (Abcam ab4680). SAG1 (DG52) and SRS9 antibodies were gifted by John Boothroyd and used as previously described^81,82^. Briefly, 40μm thick sagittal brain sections were generated, stained, and mounted on slides. Images were acquired on a Zeiss LSM 880 inverted confocal microscope (University of Arizona, Imaging Core), images were analyzed using Zen 2.6 blue edition software and counted using ImageJ software.

For quantification of cyst size in WT and *Stat1^ΔNEU^*mice, perfused brains were frozen in O.C.T Compound and 10μm sections were cut on a vibratome. Sections were mounted with ProLong Diamond Anti-Fade Mountant with DAPI according to manufacturer’s protocol. Sections were imaged on a Leica SP5-II at 60x magnification. Cysts were defined as vacuoles with >32 parasites present. Cyst area was quantified utilizing Imaris microscopy imaging software and images were generated with LAS AF software. 5-20 vacuoles per brain were quantified.

### Quantification of serum IFN-γ

Blood was collected by submandibular bleed and clotted at room temperature for 1 hour. Serum was separated through centrifugation for 10 minutes at 14,000g. Serum was diluted between 1:5 and 1:20 and IFN-γ levels were assayed by BD Mouse IFN-γ Flex Set Cytometric Bead Array according to manufacturer’s protocol. Beads were analyzed by flow cytometry on a BD Canto.

### Quantification of parasites by PCR

Sagittal sections of brain tissue were snap frozen and stored at -20°C. DNA was isolated using QIAGEN DNeasy Blood and Tissue Kit according to manufacturer’s protocol. DNA quality and concentration was assessed on a Nanodrop ND-1000 UV-Vis Spectrophotometer. 200ng of DNA was used to quantify parasite burden by qPCR with Power SYBR Green Master Mix and *T. gondii* specific primers: (forward) 5’-TCCCCTCTGCTGGCGAAAAGT-3’ and (reverse) 5’-AGCGTTCGTGGTCAACTATCGATT G-3’. qPCR was performed on Applied Biosystems ViiA7 with the following conditions: Hold phase 2min 50C, 10min 95C; PCR phase (occurs 50x) 15s at 95C, 1min @60C.

### Statistical information

Statistical tests were run in Prism software (Graphpad). Data were analyzed by two-sided student’s T-tests, two-sided student’s T-tests with Bonferroni-Dunn correction for multiple tests, 2-way analysis of variance (ANOVA), or Mann-Whitney tests where indicated. Not significant p≥0.05; *p<0.05; **p<0.01; ***p<0.001; ****p<0.0001.

## Data and Code availability

All code relating to the generation of ODEs will be made available on github.

## Acknowledgements

We thank Anthony T. Phan for assistance with experiments.

## Author contributions

Conceptualization: LAS, CAH, AAK, SL

Methodology: LAS, AW, JNE, SC, CJG, EFM, FD

Investigation: LAS, JNE, AW, SC, CJG, EFM, EW, DAC, DL, MB, MJ

Funding acquisition: LAS, CAH, AAK, EK

Supervision: LAS, CAH, AAK, SL, FD, EK

Writing – original draft: LAS, CAH

Writing – review & editing: CAH, JNE, AW, AAK, SL, SC, EW, DAC, MB, LAS

## Materials and Correspondence

Christopher Hunter;

---

**Extended Data Fig. 1:**
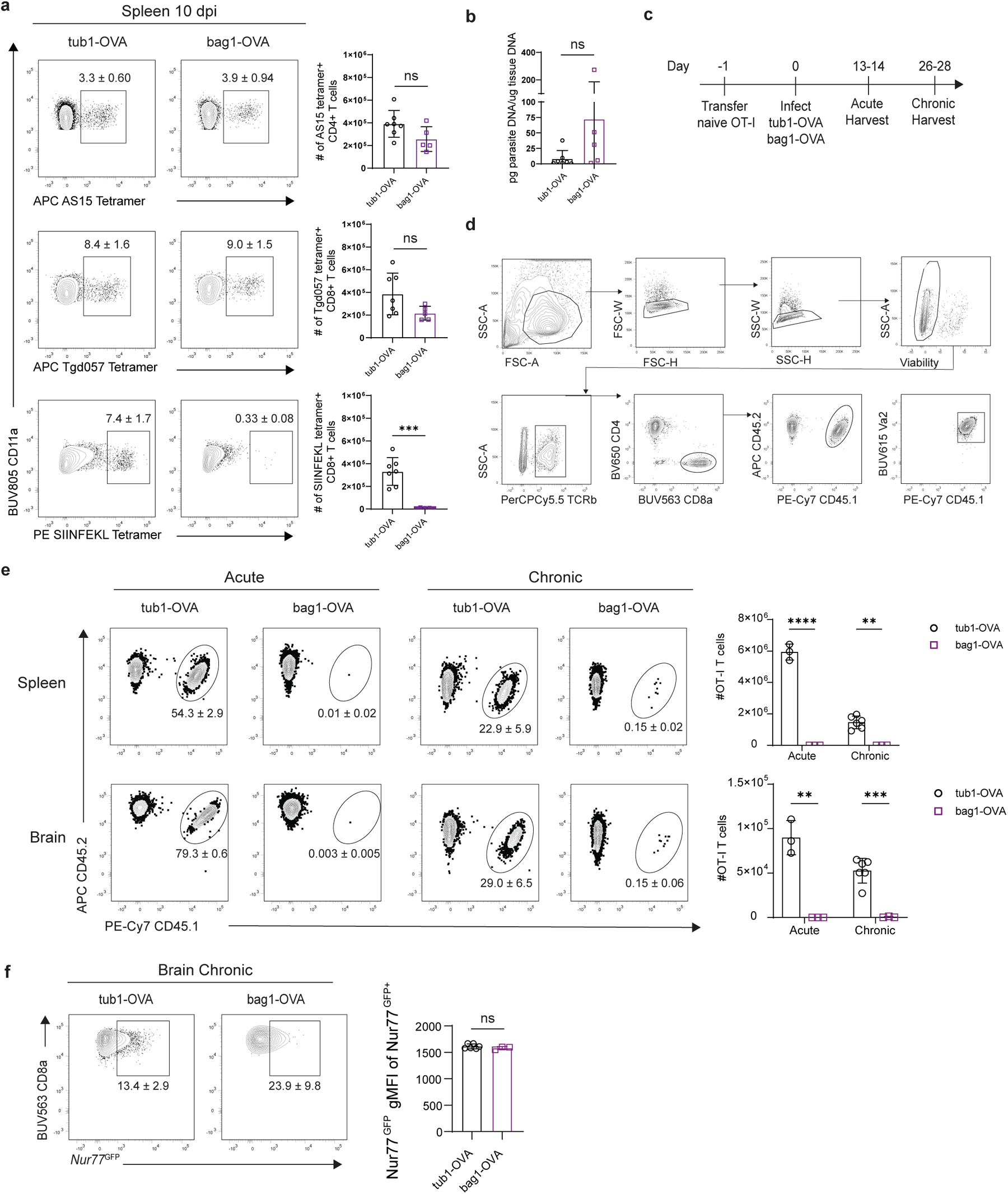
Virulence and T cell responses during early stages of infection with tub1-OVA and bag1-OVA parasites. **a**, Frequency and number of *T. gondii* peptide and SIINFEKL tetramer^+^ CD4^+^ and CD8^+^ T cells from the spleens of mice 10 dpi with tub1-OVA (left, open circles) or bag1-OVA (right, open squares) parasites. **b**, Quantification of splenic parasite burden by qPCR at 10 dpi. **c**, Experimental design for **d-f**. Naive CD45.1^+^CD45.2^+^ Nur77^GFP^ OT-I T cells were transferred 1 day prior to infection with tub1-OVA or bag1-OVA parasites. Brains and spleens were harvested from infected mice and analyzed by flow cytometry during acute and chronic infection. **d**, Gating strategy for identifying OT-I T cells by flow cytometry. **e**, Frequency and number of OT-I T cells in the spleen and brain of tub1-OVA or bag1-OVA infected mice during acute and chronic infection. **f**, Frequency and gMFI of Nur77^GFP^ expression in brain OT-I T cells following tub1-OVA or bag1-OVA infection. Data are representative of 2 independent experiments with 3-7 mice per group. Bar graphs depict the mean ± SD. Data analyzed by two-tailed unpaired Student’s *t*-test; ns *p* > 0.05, \**p* < 0.05, \*\**p* < 0.01, \*\*\**p* < 0.001, and \*\*\*\**p* < 0.0001.

**Extended Data Fig. 2:**
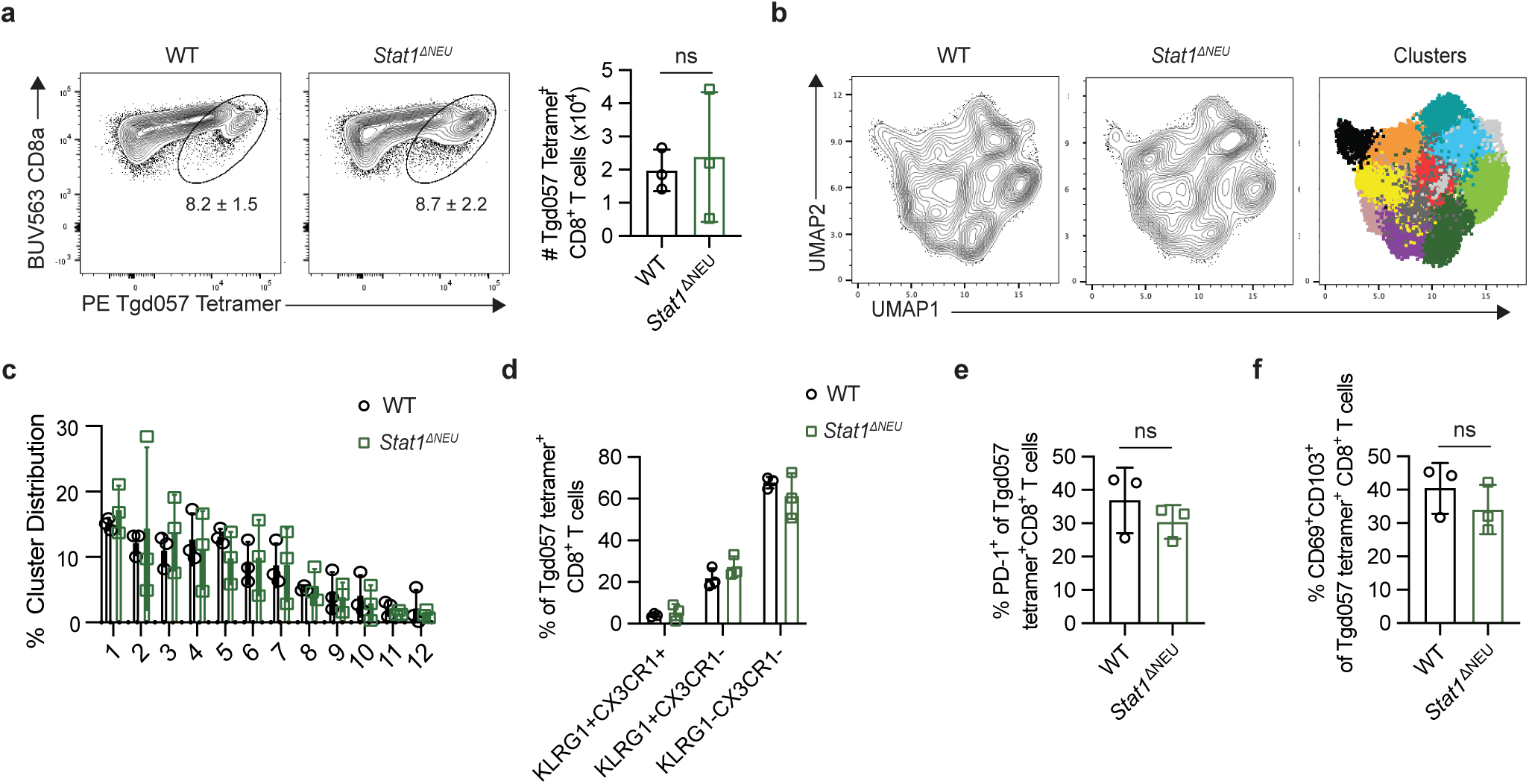
Similar *T. gondii* specific T cell responses are induced in the brains of WT and *Stat1^ΔNEU^* mice at 3 months post-infection. The brains from WT and *Stat1^ΔNEU^* mice were harvested and analyzed at 3 mpi with *T. gondii. T. gondii-*specific T cells were quantified and phenotyped by flow cytometry. **a**, Frequency and number of *T. gondii* tetramer^+^ CD8^+^ T cells. **b**, UMAP and X-shift unsupervised clustering analysis of tetramer^+^ T cells. Colors represent 12 individual clusters. **c**, Distribution of tetramer^+^ T cells across the clusters identified in **b**. **d**-**f**, Phenotyping of tetramer^+^ CD8^+^ T cells by flow cytometry. Data are representative of 2 independent experiments with 3 mice per group. Bar graphs indicate the mean ± SD. Data analyzed by two-tailed unpaired Student’s *t*-test; Bonferroni-Dunn correction for multiple comparisons included in statistical analyses performed in **c** and **d**; ns *p* > 0.5.

**Extended Data Fig. 3:**
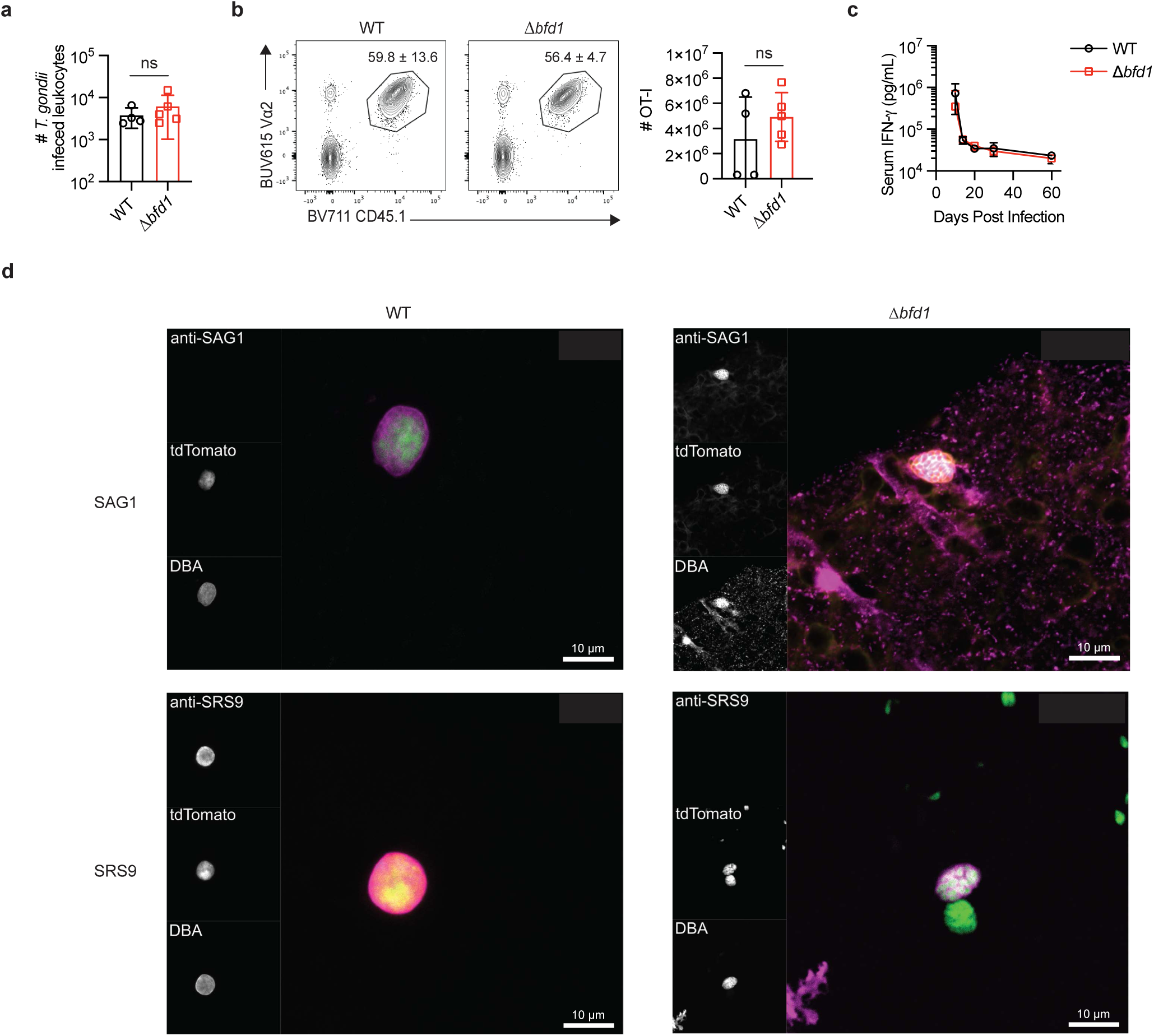
WT and *Δbfd1* parasites induce comparable immune responses during acute infection. **a**, Quantification of *T. gondii* infected cells by flow cytometry in the spleens of mice 14 dpi with tdTomato^+^ WT (black, open circles) or *Δbfd1* (red, open squares) parasites. **b**, Naive CD45.1^+^CD45.2^+^ OT-I T cells were transferred one day prior to infection with WT or *Δbfd1* parasites. OT-I T cell frequency and numbers from the spleens of mice at 14 dpi. **c**, Quantification of serum IFN-γ levels during the course of infection. **d**, Representative images of *T. gondii* in the brains of mice 25 dpi with WT (left) or *Δbfd1* (right) parasites. Brain sections were stained with DBA and either anti-SAG1 antibody (top row) or anti-SRS9 antibody (bottom row). SAG1 or SRS9 (red); tdTomato (green); DBA (purple). Scale bar = 10 μm. Data are representative of 2 independent experiments with 3-5 mice per group. Bar graphs depict the mean ± SD. Data analyzed by two-tailed unpaired Student’s *t-*test; ns *p* >0.5.

**Extended Data Fig. 4:**
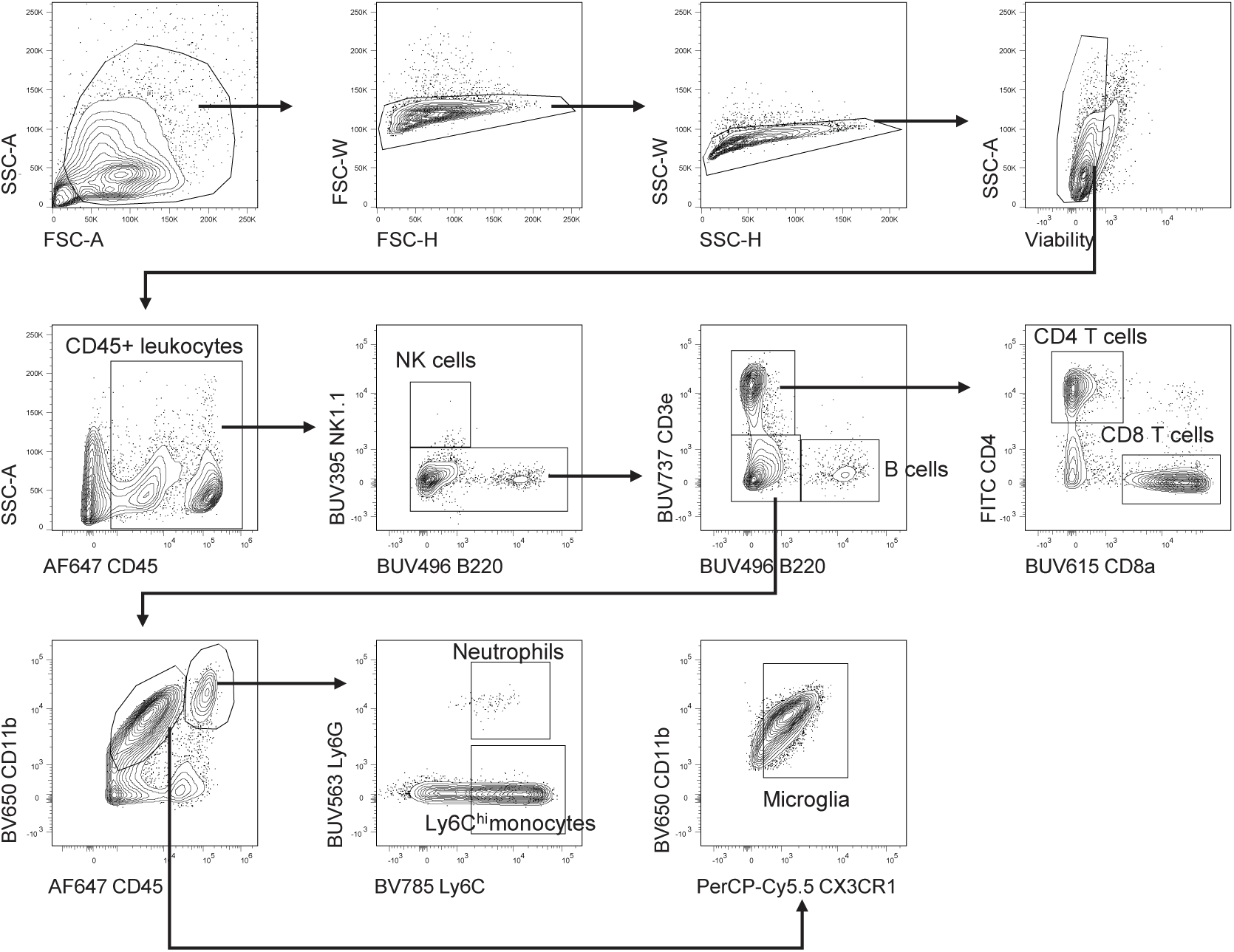
Flow cytometry gating for immune cell populations in the CNS. **a**, Flow cytometry gating strategy for CNS immune cell populations analyzed in Fig. 5 and Extended Data Figs. 5 and 6

**Extended Data Fig. 5:**
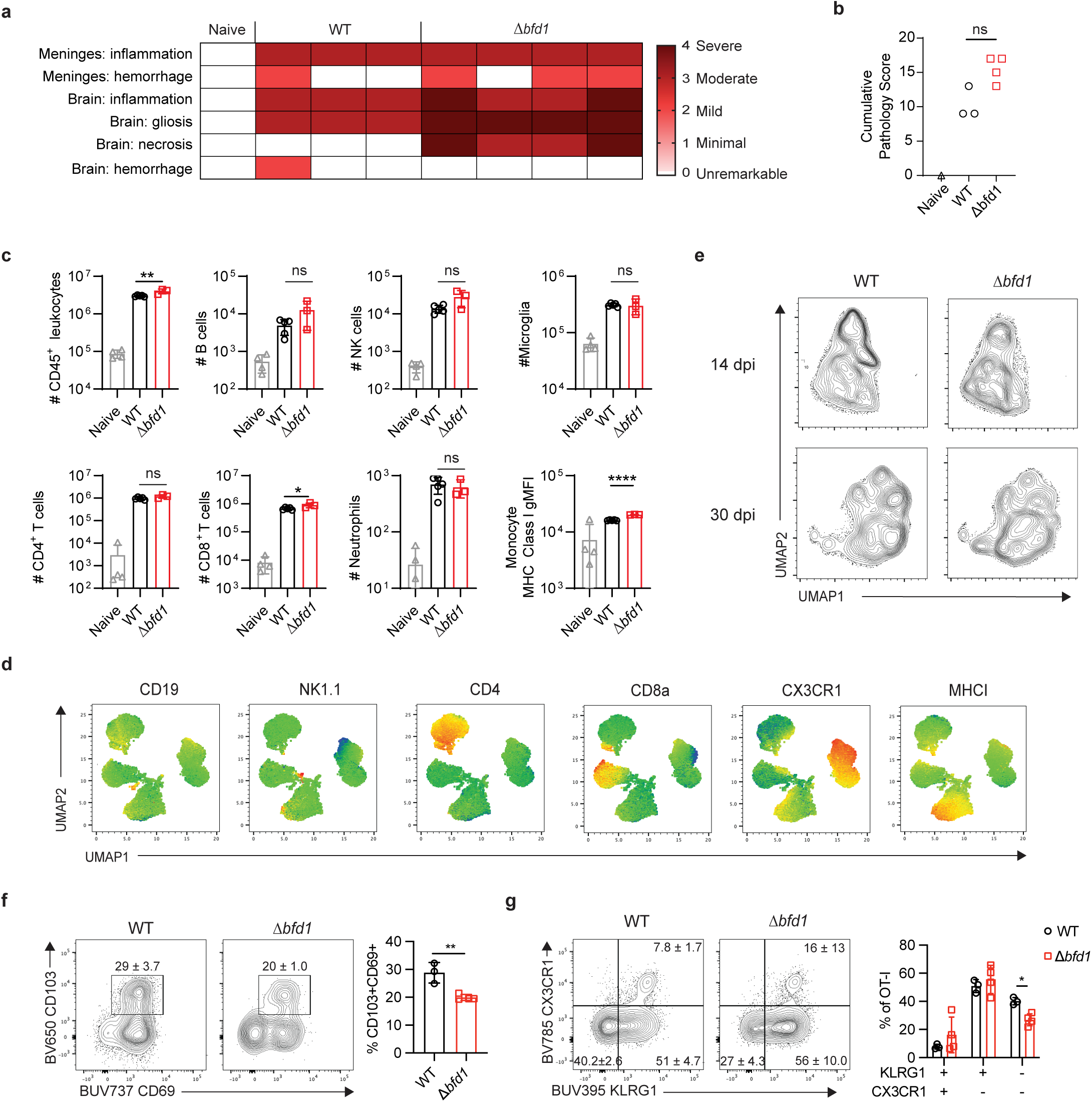
Immune responses and pathology in the brain during chronic WT and ***Δbfd1*** infection. Brains from naïve mice (grey, open triangles) and mice 30 dpi with WT (black, open circles) or *Δbfd1* (red, open squares) parasites were harvested and analyzed. **a**, Detailed pathological scoring of brain histology sections. **b**, Cumulative pathology score based on the parameters shown in **a**. **c, d**, Flow cytometry analysis of naïve brains and brains at 30 dpi. **c**, Quantification of CNS immune populations. **d**, Heatmaps displaying MFI of phenotypic markers used to identify UMAP clusters in Fig. 5e,**f**. **e**-**g**, Naive CD45.1^+^CD45.2^+^ OT-I T cells were transferred i.v. into mice 1 day prior to infection with WT (black) or *Δbfd1* (red) parasites. **e**, UMAP analysis of OT-I T cells in the brain at 14 and 30 dpi. **f**, Frequency of CD69^+^CD103^+^ OT-I T cells in the brain at 30 dpi. **g**, KLRG1 and CX3CR1 expression on OT-I T cells in the brain at 30 dpi. Data are representative of 2-3 independent experiments with 3-5 mice per group. Bar graphs depict the mean ± SD. Data analyzed by (**b**) Mann-Whitney test, (**c, f**) unpaired Student’s *t*-test; Bonferroni-Dunn correction for multiple comparisons was included in **g**; ns *p* > 0.05, \**p* < 0.05, \*\**p* < 0.01, \*\*\**p* < 0.001, and \*\*\*\**p* < 0.0001.

**Extended Data Fig. 6:**
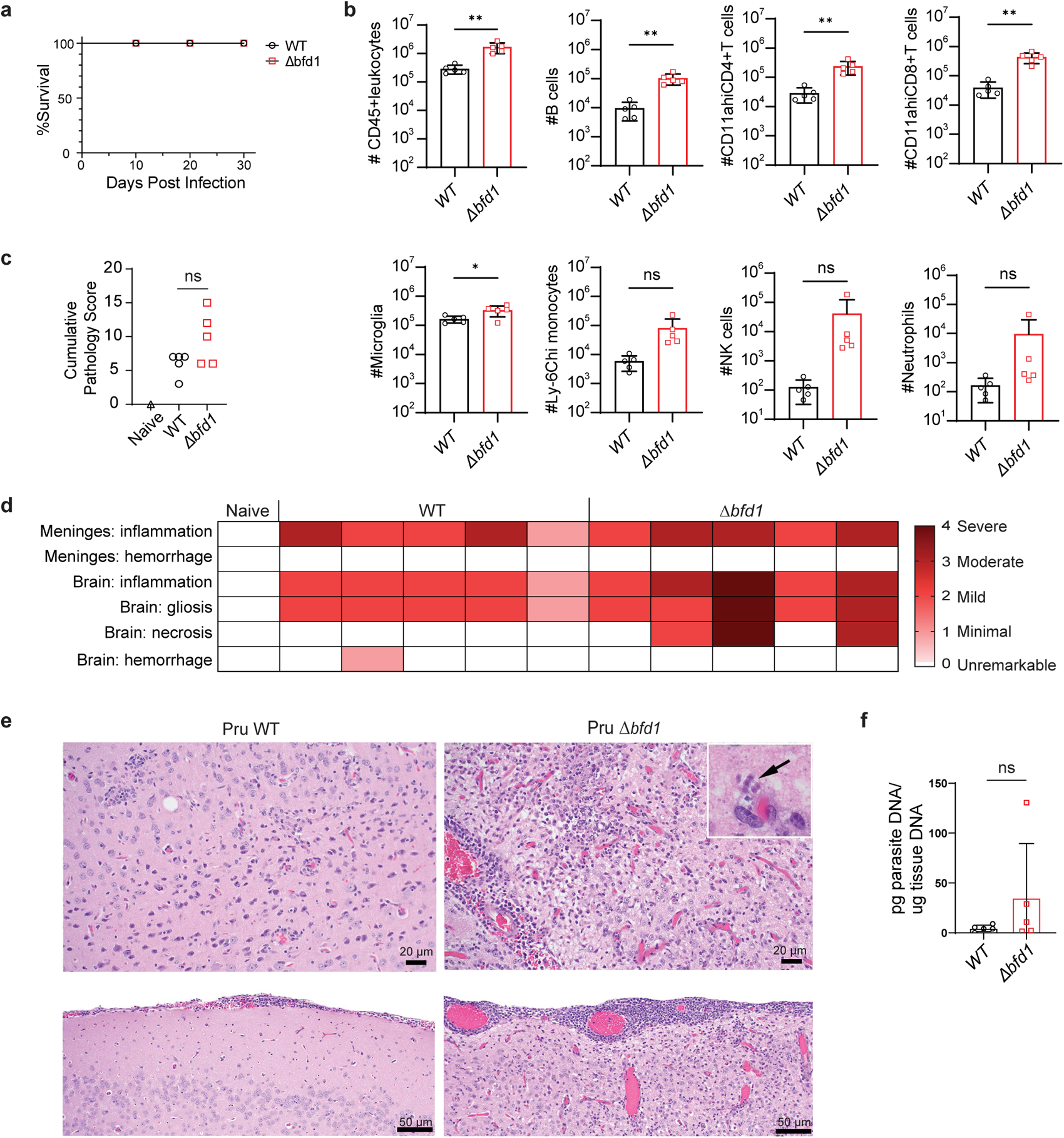
CNS inflammation and pathology associated with *Δbfd1* infection across mouse and parasite strains. **a**-**f**, BALB/c mice were infected with WT or *Δbfd1* parasites. Brains were harvested and analyzed at 30 dpi. **a**, Survival of mice until time of harvest. **b**, Quantification of CNS immune cell populations measured by flow cytometry. Gating scheme to identify populations depicted in Extended Data Fig. 4. **c,d**, Cumulative pathology score (**c**) and detailed pathological assessment (**d**) of brain histology sections. Assessment was performed by a board-certified veterinary pathologist. **e**, Representative photomicrographs of the brain showing the parenchyma (top) and meninges (bottom) at 30 dpi. Inset and black arrow show tachyzoites in the brain during *Δbfd1* infection. **f**, Quantification of CNS parasite burden by qPCR. Data are representative of 1 experiment with 5 mice per group. Bar graphs depict the mean ± SD. Data analyzed by (**b, f**) two-tailed unpaired Student’s *t*-test or (**c**) Mann-Whitney test; ns *p* > 0.05, \**p* < 0.05, \*\**p* < 0.01.

## References

1 Pirofski, L.-a. & Casadevall, A. The state of latency in microbial pathogenesis. The Journal of Clinical Investigation 130, 4525–4531 (2020). 10.1172/JCI136221

2 Mackowiak, P. A. Microbial latency. Rev Infect Dis 6, 649–668 (1984). 10.1093/clinids/6.5.649

3 Munz, C. Latency and lytic replication in Epstein-Barr virus-associated oncogenesis. Nat Rev Microbiol 17, 691–700 (2019). 10.1038/s41579-019-0249-7

4 Singh, N. & Tscharke, D. C. Herpes Simplex Virus Latency Is Noisier the Closer We Look. J Virol 94 (2020). 10.1128/JVI.01701-19

5 Alanio, A. Dormancy in Cryptococcus neoformans: 60 years of accumulating evidence. J Clin Invest 130, 3353–3360 (2020). 10.1172/JCI136223

6 Zhao, X. Y. & Ewald, S. E. The molecular biology and immune control of chronic Toxoplasma gondii infection. J Clin Invest 130, 3370–3380 (2020). 10.1172/JCI136226

7 Goodrum, F. Human Cytomegalovirus Latency: Approaching the Gordian Knot. Annu Rev Virol 3, 333–357 (2016). 10.1146/annurev-virology-110615-042422

8 Schäfer, C., Zanghi, G., Vaughan, A. M. & Kappe, S. H. I. Plasmodium vivax Latent Liver Stage Infection and Relapse: Biological Insights and New Experimental Tools. Annual Review of Microbiology 75, 87–106 (2021). 10.1146/annurev-micro-032421-061155

9 Sokol-Borrelli, S. L., Coombs, R. S. & Boyle, J. P. A Comparison of Stage Conversion in the Coccidian Apicomplexans Toxoplasma gondii, Hammondia hammondi, and Neospora caninum. Front Cell Infect Microbiol 10, 608283 (2020). 10.3389/fcimb.2020.608283

10 Matta, S. K., Rinkenberger, N., Dunay, I. R. & Sibley, L. D. Toxoplasma gondii infection and its implications within the central nervous system. Nat Rev Microbiol 19, 467–480 (2021). 10.1038/s41579-021-00518-7

11 Cerutti, A., Blanchard, N. & Besteiro, S. The Bradyzoite: A Key Developmental Stage for the Persistence and Pathogenesis of Toxoplasmosis. Pathogens 9 (2020). 10.3390/pathogens9030234

12. Waldman, B. S. et al. Identification of a Master Regulator of Differentiation in Toxoplasma.

13 Licon, M. H. et al. A positive feedback loop controls Toxoplasma chronic differentiation. Nat Microbiol 8, 889–904 (2023). 10.1038/s41564-023-01358-2

14. Sokol-Borrelli, S. L. et al. A transcriptional network required for bradyzoite development in Toxoplasma gondii is dispensable for recrudescent disease.

15 Su, C. et al. Recent expansion of Toxoplasma through enhanced oral transmission. Science 299, 414–416 (2003). 10.1126/science.1078035

16 Coombes, J. L. & Robey, E. A. Dynamic imaging of host-pathogen interactions in vivo. Nat Rev Immunol 10, 353–364 (2010). 10.1038/nri2746

17 John, B., Weninger, W. & Hunter, C. A. Advances in imaging the innate and adaptive immune response to Toxoplasma gondii. Future Microbiol 5, 1321–1328 (2010). 10.2217/fmb.10.97

18 Watts, E. et al. Novel Approaches Reveal that Toxoplasma gondii Bradyzoites within Tissue Cysts Are Dynamic and Replicating Entities In Vivo. mBio 6, e01155–01115 (2015). 10.1128/mBio.01155-15

19 Tomita, T. et al. Toxoplasma gondii Matrix Antigen 1 Is a Secreted Immunomodulatory Effector. mBio 12 (2021). 10.1128/mBio.00603-21

20 Mayoral, J., Shamamian, P., Jr. & Weiss, L. M. In Vitro Characterization of Protein Effector Export in the Bradyzoite Stage of Toxoplasma gondii. mBio 11 (2020). 10.1128/mBio.00046-20

21 Burke, J. M., Roberts, C. W., Hunter, C. A., Murray, M. & Alexander, J. Temporal differences in the expression of mRNA for IL-10 and IFN-gamma in the brains and spleens of C57BL/10 mice infected with Toxoplasma gondii. Parasite Immunol 16, 305–314 (1994).

22 Young, J. D. & McGwire, B. S. Infliximab and reactivation of cerebral toxoplasmosis. N Engl J Med 353, 1530–1531; discussion 1530-1531 (2005). 10.1056/NEJMc051556

23 Luft, B. J. & Remington, J. S. Toxoplasmic Encephalitis. The Journal of Infectious Diseases 157, 1–6 (1988).

24 Gazzinelli, R. T. et al. In the absence of endogenous IL-10, mice acutely infected with Toxoplasma gondii succumb to a lethal immune response dependent on CD4+ T cells and accompanied by overproduction of IL-12, IFN-gamma and TNF-alpha. J Immunol 157, 798–805 (1996).

25 Villarino, A. et al. The IL-27R (WSX-1) is required to suppress T cell hyperactivity during infection. Immunity 19, 645–655 (2003).

26 Wilson, E. H., Wille-Reece, U., Dzierszinski, F. & Hunter, C. A. A critical role for IL-10 in limiting inflammation during toxoplasmic encephalitis. J Neuroimmunol 165, 63–74 (2005). 10.1016/j.jneuroim.2005.04.018

27 Deckert-Schluter, M. et al. Interleukin-10 downregulates the intracerebral immune response in chronic Toxoplasma encephalitis. Journal Neuroimmunology 76, 167–176 (1997).

28 Stumhofer, J. S. et al. Interleukin 27 negatively regulates the development of interleukin 17-producing T helper cells during chronic inflammation of the central nervous system. Nat Immunol 7, 937–945 (2006). 10.1038/ni1376

29 Aliberti, J. Host persistence: exploitation of anti-inflammatory pathways by Toxoplasma gondii. Nat Rev Immunol 5, 162–170 (2005).

30 Buckley, M. W. & McGavern, D. B. Immune dynamics in the CNS and its barriers during homeostasis and disease. Immunol Rev 306, 58–75 (2022). 10.1111/imr.13066

31 Klein, R. S. & Hunter, C. A. Protective and Pathological Immunity during Central Nervous System Infections. Immunity 46, 891–909 (2017). 10.1016/j.immuni.2017.06.012

32 Miller, K. D., Schnell, M. J. & Rall, G. F. Keeping it in check: chronic viral infection and antiviral immunity in the brain. Nat Rev Neurosci 17, 766–776 (2016). 10.1038/nrn.2016.140

33 Rose, R. W., Vorobyeva, A. G., Skipworth, J. D., Nicolas, E. & Rall, G. F. Altered levels of STAT1 and STAT3 influence the neuronal response to interferon gamma. J Neuroimmunol 192, 145–156 (2007). 10.1016/j.jneuroim.2007.10.007

34 Chandrasekaran, S. et al. IFN-gamma stimulated murine and human neurons mount anti-parasitic defenses against the intracellular parasite Toxoplasma gondii. Nat Commun 13, 4605 (2022). 10.1038/s41467-022-32225-z

35 Chevalier, G. et al. Neurons are MHC class I-dependent targets for CD8 T cells upon neurotropic viral infection. PLoS Pathog 7, e1002393 (2011). 10.1371/journal.ppat.1002393

36. 36 Moseman, E. A., Blanchard, A. C., Nayak, D. & McGavern, D. B. T cell engagement of cross-presenting microglia protects the brain from a nasal virus infection. Sci Immunol 5 (2020). 10.1126/sciimmunol.abb1817

37 Binder, G. K. & Griffin, D. E. Interferon-gamma-mediated site-specific clearance of alphavirus from CNS neurons. Science 293, 303–306 (2001). 10.1126/science.1059742

38 Patterson, C. E., Lawrence, D. M., Echols, L. A. & Rall, G. F. Immune-mediated protection from measles virus-induced central nervous system disease is noncytolytic and gamma interferon dependent. J Virol 76, 4497–4506 (2002).

39 Burdeinick-Kerr, R., Govindarajan, D. & Griffin, D. E. Noncytolytic clearance of sindbis virus infection from neurons by gamma interferon is dependent on Jak/STAT signaling. J Virol 83, 3429–3435 (2009). 10.1128/JVI.02381-08

40 Schluter, D., Deckert, M., Hof, H. & Frei, K. Toxoplasma gondii infection of neurons induces neuronal cytokine and chemokine production, but gamma interferon- and tumor necrosis factor-stimulated neurons fail to inhibit the invasion and growth of T. gondii. Infect Immun 69, 7889–7893 (2001).

41 Denkers, E. Y. et al. Perforin-mediated cytolysis plays a limited role in host resistance to Toxoplasma gondii. Journal of immunology 159, 1903–1908 (1997).

42 Suzuki, Y. et al. Removal of Toxoplasma gondii cysts from the brain by perforin-mediated activity of CD8+ T cells. Am J Pathol 176, 1607–1613 (2010). 10.2353/ajpath.2010.090825

43 Salvioni, A. et al. Robust Control of a Brain-Persisting Parasite through MHC I Presentation by Infected Neurons. Cell Rep 27, 3254–3268 e3258 (2019). 10.1016/j.celrep.2019.05.051

44 Suzuki, Y., Orellana, M. A., Schreiber, R. D. & Remington, J. S. Interferon-gamma: the major mediator of resistance against Toxoplasma gondii. Science 240, 516–518 (1988). 10.1126/science.3128869

45 Wilson, E. H. et al. Behavior of parasite-specific effector CD8+ T cells in the brain and visualization of a kinesis-associated system of reticular fibers. Immunity 30, 300–311 (2009).

46 Schaeffer, M. et al. Dynamic imaging of T cell-parasite interactions in the brains of mice chronically infected with Toxoplasma gondii. J Immunol 182, 6379–6393 (2009). 10.4049/jimmunol.0804307

47 Shallberg, L. A. et al. Impact of secondary TCR engagement on the heterogeneity of pathogen-specific CD8+ T cell response during acute and chronic toxoplasmosis. PLoS Pathog 18, e1010296 (2022). 10.1371/journal.ppat.1010296

48 John, B. et al. Analysis of behavior and trafficking of dendritic cells within the brain during toxoplasmic encephalitis. PLoS Pathog 7, e1002246 (2011).

49 Tiwari, A. et al. Penetration of CD8(+) Cytotoxic T Cells into Large Target, Tissue Cysts of Toxoplasma gondii, Leads to Its Elimination. Am J Pathol 189, 1594–1607 (2019). 10.1016/j.ajpath.2019.04.018

50 Fischer, H. G., Bonifas, U. & Reichmann, G. Phenotype and functions of brain dendritic cells emerging during chronic infection of mice with *Toxoplasma gondii*. Journal Immunology 164, 4826–4834 (2000).

51 Kim, S. K., Karasov, A. & Boothroyd, J. C. Bradyzoite-specific surface antigen SRS9 plays a role in maintaining Toxoplasma gondii persistence in the brain and in host control of parasite replication in the intestine. Infect Immun 75, 1626–1634 (2007). 10.1128/iai.01862-06

52 Sullivan, A. et al. A mathematical model for within-host Toxoplasma gondii invasion dynamics. Math Biosci Eng 9, 647–662 (2012). 10.3934/mbe.2012.9.647

53 Sullivan, A. M. et al. Evidence for finely-regulated asynchronous growth of Toxoplasma gondii cysts based on data-driven model selection. PLoS Comput Biol 9, e1003283 (2013). 10.1371/journal.pcbi.1003283

54 Harris, T. H. et al. Generalized Levy walks and the role of chemokines in migration of effector CD8+ T cells. Nature 486, 545–548 (2012).

55. Christian, D. A., et al. cDC1 coordinate innate and adaptive responses in the omentum required for T cell priming and memory. Sci Immunol 7, eabq7432 (2022). 10.1126/sciimmunol.abq7432

56 Baez, J. C., Biamonte, J.D. Quantum techniques in stochastic mechanics. (World Scientific, 2018).

57 Yap, G. S. & Sher, A. Effector Cells of Both Nonhemopoietic and Hemopoietic Origin Are Required for Interferon (IFN)-γ– and Tumor Necrosis Factor (TNF)-α–dependent Host Resistance to the Intracellular Pathogen, Toxoplasma gondii. Journal of Experimental Medicine 189, 1083–1092 (1999). 10.1084/jem.189.7.1083

58 Suzuki, Y., Conley, F. K. & Remington, J. S. Importance of endogenous IFN-gamma for prevention of toxoplasmic encephalitis in mice. J Immunol 143, 2045–2050 (1989).

59 Suzuki, Y., Joh, K., Orellana, M. A., Conley, F. K. & Remington, J. S. A gene(s) within the H-2D region determines the development of toxoplasmic encephalitis in mice. Immunology 74, 732–739 (1991).

60 Bergersen, K. V., Barnes, A., Worth, D., David, C. & Wilson, E. H. Targeted Transcriptomic Analysis of C57BL/6 and BALB/c Mice During Progressive Chronic Toxoplasma gondii Infection Reveals Changes in Host and Parasite Gene Expression Relating to Neuropathology and Resolution. Front Cell Infect Microbiol 11, 645778 (2021). 10.3389/fcimb.2021.645778

61 Rall, G. F., Mucke, L. & Oldstone, M. B. Consequences of cytotoxic T lymphocyte interaction with major histocompatibility complex class I-expressing neurons in vivo. Journal of Experimental Medicine 182, 1201–1212 (1995). 10.1084/jem.182.5.1201

62 Gazzinelli, R., Xu, Y., Hieny, S., Cheever, A. & Sher, A. Simultaneous depletion of CD4+ and CD8+ T lymphocytes is required to reactivate chronic infection with Toxoplasma gondii. J Immunol 149, 175–180 (1992).

63 Ma, J. S. et al. Selective and strain-specific NFAT4 activation by the Toxoplasma gondii polymorphic dense granule protein GRA6. J Exp Med 211, 2013–2032 (2014). 10.1084/jem.20131272

64 Sa, Q. et al. Determination of a Key Antigen for Immunological Intervention To Target the Latent Stage of Toxoplasma gondii. J Immunol 198, 4425–4434 (2017). 10.4049/jimmunol.1700062

65 Herz, J., Johnson, K. R. & McGavern, D. B. Therapeutic antiviral T cells noncytopathically clear persistently infected microglia after conversion into antigen-presenting cells. J Exp Med 212, 1153–1169 (2015). 10.1084/jem.20142047

66 Cabral, C. M. et al. Neurons are the Primary Target Cell for the Brain-Tropic Intracellular Parasite Toxoplasma gondii. PLoS Pathog 12, e1005447 (2016). 10.1371/journal.ppat.1005447

67 Frickel, E. M. & Hunter, C. A. Lessons from Toxoplasma: Host responses that mediate parasite control and the microbial effectors that subvert them. J Exp Med 218 (2021). 10.1084/jem.20201314

68. Gay, G. A.-O. et al. Toxoplasma gondii TgIST co-opts host chromatin repressors dampening STAT1-dependent gene regulation and IFN-γ-mediated host defenses.

69 Seizova, S. et al. Transcriptional modification of host cells harboring Toxoplasma gondii bradyzoites prevents IFN gamma-mediated cell death. Cell Host Microbe 30, 232–247.e236 (2022). 10.1016/j.chom.2021.11.012

70. Olias, P., Etheridge, R. D., Zhang, Y., Holtzman, M. J. & Sibley, L. D. Toxoplasma Effector Recruits the Mi-2/NuRD Complex to Repress STAT1 Transcription and Block IFN-γ-Dependent Gene Expression.

71 Hidano, S. et al. STAT1 Signaling in Astrocytes Is Essential for Control of Infection in the Central Nervous System. mBio 7 (2016). 10.1128/mBio.01881-16

72 Landrith, T. A. et al. CD103(+) CD8 T Cells in the Toxoplasma-Infected Brain Exhibit a Tissue-Resident Memory Transcriptional Profile. Front Immunol 8, 335 (2017). 10.3389/fimmu.2017.00335

73 Bhadra, R., Gigley, J. P., Weiss, L. M. & Khan, I. A. Control of Toxoplasma reactivation by rescue of dysfunctional CD8+ T-cell response via PD-1-PDL-1 blockade. Proc Natl Acad Sci U S A (2011).

74 Chauhan, P. & Lokensgard, J. R. Glial Cell Expression of PD-L1. Int J Mol Sci 20 (2019). 10.3390/ijms20071677

75 Meerschaert, K. A. et al. Neuronally expressed PDL1, not PD1, suppresses acute nociception. Brain, Behavior, and Immunity 106, 233–246 (2022). 10.1016/j.bbi.2022.09.001

76 Handel, A., La Gruta, N. L. & Thomas, P. G. Simulation modelling for immunologists. Nature Reviews Immunology 20, 186–195 (2020). 10.1038/s41577-019-0235-3

77 Kirschner, D., Pienaar, E., Marino, S. & Linderman, J. J. A review of computational and mathematical modeling contributions to our understanding of Mycobacterium tuberculosis within-host infection and treatment. Curr Opin Syst Biol 3, 170–185 (2017). 10.1016/j.coisb.2017.05.014

78 Yap, G. S. Avirulence: an essential feature of the parasitic lifestyle. Trends Parasitol 38, 1028–1030 (2022). 10.1016/j.pt.2022.09.003

79 Matthews, K. R., McCulloch, R. & Morrison, L. J. The within-host dynamics of African trypanosome infections. Philos Trans R Soc Lond B Biol Sci 370 (2015). 10.1098/rstb.2014.0288

80 Brunet, L. R., Finkelman, F. D., Cheever, A. W., Kopf, M. A. & Pearce, E. J. IL-4 protects against TNF-a-mediated cachexia and death during acute Schistosomiasis. Journal Immunology 159, 777–785 (1997).

81 Burg, J. L., Perelman, D., Kasper, L. H., Ware, P. L. & Boothroyd, J. C. Molecular analysis of the gene encoding the major surface antigen of Toxoplasma gondii. J Immunol 141, 3584–3591 (1988).

82 Kim, S. K. & Boothroyd, J. C. Stage-specific expression of surface antigens by Toxoplasma gondii as a mechanism to facilitate parasite persistence. J Immunol 174, 8038–8048 (2005). 10.4049/jimmunol.174.12.8038

