## Supplementary Text for "The latent stage of *Toxoplasma gondii* is targeted by the immune response and host protective"

**Supplementary Text:** Development of model for *T. gondii* infection dynamics in the CNS.

### 1. Mathematical Model

In order to model the spread of *T. gondii* within the organism, we wish to write down a system of ODEs to determine the number of tachyzoite-infected cells and bradyzoite-infected cells. In order to mathematically describe the mechanisms controlling the numbers of infected cells, it will help us to first write down a larger system of ODEs describing the dynamics of tachyzoite-infected cells $I_T$, bradyzoite-infected cells $I_B$, uninfected cells $S$, immune cells $Z$, free tachyzoites $P_T$ and free bradyzoites $P_B$. With appropriate approximations, simplified dynamics governing only the infected cells and immune cells will be derived. Throughout the derivation, the similarities and differences between our work and that of Sullivan, et al. [1] will be described.

Before parasites enter the organism, the number of cells is at an equilibrium value $S_0$. The natural death rate of healthy cells is $d_S$. When a free parasite in a particular phase encounters a target cell, an infection occurs, so the number of free parasites of that phase and the number of healthy cells is reduced by one while the number of infected cells of that phase increases by one. Suppose the parasite-induced death rate (burst rate) of the tachyzoite-infected cells is $d_T$, and $n_T$ parasites are released on average. The increase in $P_T$ is then proportional to $I_T$ with growth rate $d_T n_T$. For a given free parasite, if the probability per unit time of encountering a cell is $\beta_{PT}N/S_0$ (written in this peculiar way to highlight the fact that the total contact rate increases as the cell population increases) and the probability of that cell being a susceptible cell is $S/N$, then the rate at which a single parasite finds a new host is $\beta_{PT}S/S_0$. The clearance rate of the free parasite (due to means other than entering a healthy cell) is given by $u_T$. Tachyzoite-infected cells can also die if targeted by immune cells $Z$ which have a natural decay rate of $\mu$. The tachyzoite-induced immune activation rate is $a_T$, which induces a tachyzoite-infected cell clearance rate of $\psi_T$. All of these definitions can be applied analogously for the bradyzoite-infected cells and free parasites, but note that any free parasite re-enters the cell as a tachyzoite. (Equivalently, as soon as the free bradyzoites enter a new host cell, there is a rapid transition to the tachyzoite phase). The one additional consideration is the differentiation rate $c_{TB}$ from tachyzoites to bradyzoites. The result is the following set of coupled nonlinear ODEs.

$$\frac{dS}{dt} = -\beta_{PT}\frac{SP_T}{S_0} - \beta_{PB}\frac{SP_B}{S_0} + d_S(S_0 - S) \tag{1}$$

$$\frac{dI_T}{dt} = +\beta_{PT}\frac{SP_T}{S_0} + \beta_{PB}\frac{SP_B}{S_0} - d_T I_T - \psi_T\frac{I_T Z}{S_0} - c_{TB}I_T \tag{2}$$

$$\frac{dI_B}{dt} = -d_B I_B - \psi_B\frac{I_B Z}{S_0} + c_{TB}I_T \tag{3}$$

$$\frac{dP_T}{dt} = -\beta_{PT}\frac{SP_T}{S_0} + d_T n_T I_T - u_T P_T \tag{4}$$

$$\frac{dP_B}{dt} = -\beta_{PB}\frac{SP_B}{S_0} + d_B n_B I_B - u_B P_B \tag{5}$$

$$\frac{dZ}{dt} = a_T\frac{I_T Z}{S_0} + a_B\frac{I_B Z}{S_0} - \mu Z \tag{6}$$

Up to naming conventions, this system closely resembles equations (1) through (7) of [1], but there are some notable differences. They separately treat the dynamics of cells containing early-stage bradyzoites and encysted bradyzoites; here, we combine these cells into a single parameter $I_B$. They do not consider the possibility of immune activation and clearance of bradyzoites, we do which allows us to assess the possible impact of an immune-mediated mechanism to limit cyst numbers. They consider a more complicated immune activation rate that saturates when the number of tachyzoite-infected cells is large, whereas our immune activation is simply linear in $I_T$. We do not feel this is necessary since the negative feedback between infected cells and immune cells can be studied with our simpler model.

The dependence on the free parasites can be eliminated by making a quasi-steady state approximation. The fundamental assumption we will be making is that the parasites are cleared much faster than the lifetime of the parasites within the cell $\beta_{PT}s + u_T \gg d_T$, where $s \equiv S/S_0$. Using $d_T$ as the characteristic time governing cell dynamics, we arrive at the following approximation:

$$\frac{dP_T}{d(d_T t)} \ll \frac{\beta_{PT} s + u_T}{d_T} P_T. \qquad (7)$$

Rather than working in terms of $\beta_{PT}$ and $u_T$, we can interpret $f_T(t) \equiv \frac{\beta_{PT} s}{\beta_{PT} s + u_T}$ as the fraction of free parasites which (eventually) go on to infect a new cell. This fraction is theoretically time-dependent because as the number of susceptible cells is reduced, more parasites die without finding a host. However, since $s$ presumably stays close to one, it is useful to define the quantity $f_{T0} \equiv \frac{\beta_{PT}}{\beta_{PT} + u_T}$ which is the fraction of free parasites which infect a new cell if all cells are susceptible (the zero stands for $t = 0$). Now, applying (7), we can drop the $\frac{dP_T}{dt}$ term in equation (4) to find that $P_T$ is proportional to $I_T$:

$$P_T = \frac{d_T n_T}{\beta_{PT} s + u_T} I_T = \frac{d_T n_T}{\beta_{PT}} \left[ \frac{f_{T0}}{f_{T0} s + (1 - f_{T0})} \right] I_T. \qquad (8)$$

We are not claiming $P_T$ is constant throughout our experiment; rather, we are saying $\frac{dP_T}{dt}$ is smaller in magnitude than the other terms in (4). The same approximation can be used to relate $P_B$ to $I_B$. It is useful to define the "contact rate" $\beta_T \equiv d_T n_T f_{T0}$ describing the rate at which tachyzoite-infected cells infect new susceptible cells when $s = 1$. In terms of these new parameters, we can re-write (1), (2), (3), and (6) under the quasi-steady state approximation. Furthermore, we will assume that $s \approx 1$ throughout the duration of the experiment such that the above quantity in brackets remains one.

$$\frac{dS}{dt} = -\beta_T I_T - \beta_B I_B + d_S(S_0 - S) \quad (9)$$

$$\frac{dI_T}{dt} = +\beta_T I_T + \beta_B I_B - d_T I_T - \psi_T \frac{I_T Z}{S_0} - c_{TB} I_T \quad (10)$$

$$\frac{dI_B}{dt} = -d_B I_B - \psi_B \frac{I_B Z}{S_0} + c_{TB} I_T \quad (11)$$

$$\frac{dZ}{dt} = a_T \frac{I_T Z}{S_0} + a_B \frac{I_B Z}{S_0} - \mu Z \quad (12)$$

This is similar to the ODEs used by Sullivan et al., equations (11) to (14), given the differences already discussed, except we assume $S$ is constant. As we showed, carefully applying a quasi-steady state approximation leads to a peculiar term, that shown in brackets, not a term proportional to $I_T\ S$ as Sullivan et al. suggest. Regardless, both models agree that when $N \approx S_0$, which we expect is the case experimentally.

Since equations (10) and (11) are now independent of s, we do not need to solve (9) unless we care to know the current number of healthy cells. However, a more useful quantity (which doesn't require us to know the natural birth/death rate) is the cumulative number of infected cells which will be denote $C$.

$$\frac{dC}{dt} = +\beta_T I_T + \beta_B I_B \tag{13}$$

Ten parameters will govern the dynamics: $\beta_T$, $\beta_B$, $d_T$, $d_B$, $c_{TB}$, $\psi_T$, $\psi_B$, $a_T$, $a_B$, $\mu$. It will always be assumed that $I_B(0) = 0$, so only three initial conditions $S(0) = S_0$, $I_T(0)$, and $Z(0)$ will need to be specified. Throughout the remaining sections, the reduced dynamics (10), (11), and (12) will be studied to determine the role played by each of the rates and explain some of the experimental findings. The system exhibits two equilibria: the parasite-free equilibrium $(I_T,I_B,Z) = (0,0,0)$ and the endemic equilibrium $(I_T,I_B,Z) = (I_T^*,I_B^*,Z^*)$. In section 2, the model will be analyzed close to the parasite-free equilibrium where the immune response is negligible to study the initial growth rates of infected cells with and without a differentiation mechanism. In section 3, the stability of the endemic equilibrium will be studied to investigate how the reactivation and immune parameters dictate the behavior at late times. In section 4, a combination of experimental data and mathematical considerations will be used to determine reasonable numerical values for the rates. Finally, in section 5, equations (10), (11), (12), and (13) will be solved to understand the mechanisms driving parasite growth, differentiation, reactivation, and immune response.

### 2. A Complete Characterization of the Linear System

At early times, the number of infected cells is small enough that the behavior is governed by the dynamics close to the parasite-free equilibrium $I_T = I_B = Z = 0$. In fact, we can ignore the immune dynamics at early times since even if $Z(0) > 0$, the leading-order response would be a gradual decay of the form $Z(t) = Z_0e^{-\mu t}$. Furthermore, since $\beta_T$ and $d_T$ only appear as a difference in our system (unless one is interested in knowing $C(t)$ which will not be considered in this section), we will work in terms of the quantity $\delta_T \equiv \beta_T - d_T$, and we are left with only four parameters and one initial condition in the linearized model: $\delta_T$, $\beta_B$, $d_T$, $c_{TB}$, $I_T(0)$.

The linearized system close to the parasite-free equilibrium reads

$$\frac{d}{dt}\begin{bmatrix} I_T \\ I_B \end{bmatrix} = \begin{bmatrix} \delta_T - c_{TB} & \beta_B \\ c_{TB} & -d_B \end{bmatrix}\begin{bmatrix} I_T \\ I_B \end{bmatrix} \quad (14)$$

The growth of the parasite is governed by the eigenvalues $\lambda_\pm$ of this matrix.

$$\begin{aligned} \lambda_\pm &= \left(\frac{\delta_T - c_{TB} - d_B}{2}\right) \pm \sqrt{\left(\frac{\delta_T - c_{TB} - d_B}{2}\right)^2 + \Big((\delta_T - c_{TB})d_B + \beta_B c_{TB}\Big)} \\ &= \left(\frac{\delta_T - c_{TB} - d_B}{2}\right) \pm \sqrt{\left(\frac{\delta_T - c_{TB} + d_B}{2}\right)^2 + \beta_B c_{TB}} \end{aligned} \quad (15)$$

The discriminant is always positive, so the eigenvalues are purely real. If at least one eigenvalue is positive, then the parasite will successfully multiply in the organism, but if both eigenvalues are negative, then the infection quickly goes away. There exists a positive eigenvalue if and only if

$$\delta_T - c_{TB} + \frac{\beta_B c_{TB}}{d_B} > 0 . \quad (16)$$

The fact that *T. gondii* is so successful at replicating in a host is proof that this equality is always satisfied.

The exact solution to (14) is

$$\begin{bmatrix} I_T(t) \\ I_B(t) \end{bmatrix} = I_T(0)\begin{bmatrix} 1 \\ 0 \end{bmatrix} e^{\lambda_- t} + \frac{I_T(0)}{\lambda_+ - \lambda_-}\begin{bmatrix} \delta_T - c_{TB} - \lambda_- \\ c_{TB} \end{bmatrix}\left(e^{\lambda_+ t} - e^{\lambda_- t}\right). \quad (17)$$

When $c_{TB} = 0$, we get that $\lambda_+ = \delta_T$ and $\lambda_- = -d_B$, the growth and decay rates for the tachyzoite- and bradyzoite-infected cells in isolation. But, since no cysts can form, the solution can be written simply as

$$\begin{bmatrix} I_T^M(t) \\ I_B^M(t) \end{bmatrix} = I_T(0) \begin{bmatrix} 1 \\ 0 \end{bmatrix} e^{\delta_T t}. \quad (18)$$

The superscript $M$ stands for mutant. So, if $c_{TB} = 0$, $\delta_T$ is the all-important quantity that sets the initial growth rate of the number of infected cells. Notably, since we see exponential growth of tachyzoites for the mutant, we know that $\delta_T > 0$. For arbitrary $c_{TB}$, we have to look at the largest eigenvalue $\lambda_+$ to determine the growth rate. Of particular interest is how $\lambda_+$ changes with $c_{TB}$.

$$\frac{d\lambda_+}{dc_{TB}} = -\frac{1}{2} + \frac{1}{2} \frac{\beta_B - \left(\frac{\delta_T - c_{TB} + d_B}{2}\right)}{\sqrt{\left(\frac{\delta_T - c_{TB} + d_B}{2}\right)^2 + \beta_B c_{TB}}} \quad (19)$$

Conveniently, $\lambda_+$ is monotonic in $c_{TB}$, meaning the condition for faster mutant growth is independent of the value of $c_{TB}$, so we might as well check the sign of the above quantity at $c_{TB} = 0$. The condition for faster mutant growth is

$$\left.\frac{d\lambda_+}{dc_{TB}}\right|_{c_{TB}=0} = -1 + \frac{\beta_B}{\delta_T + d_B} < 0 \implies \boxed{\beta_T - d_T > \beta_B - d_B}. \quad (20)$$

This is the key result of this section. At first glance, this may seem obvious: if the net growth rate of the tachyzoite-infected cells exceeds that of the bradyzoite-infected cells, then the differentiation will reduce the rate of infection spread. But, remember that $\beta_B$ is not simply the cyst growth rate; it is actually the growth rate of tachyzoite-infected cells due to the death of cysts. It's interesting that such a simple equation still holds in this case. Since the rates $\beta_T$ and $d_T$ are more well understood and $d_B$ is much smaller than either of these rates, equation (20) places a bound on the less understood rate $\beta_B$ characterizing the conversion of bradyzoites to tachyzoites. One subtlety that probably will not matter for the sake of this work is that if $\beta_B \approx \beta_T - d_T + d_B$, then (20) may give the wrong answer as to whether the mutant or wild type has faster parasite growth. In such a case, both eigenvalues may be important, and one must more carefully specify which metric we want to use to determine which infection is more severe (could use $I_T(t)$, $C(t)$, etc.).

The magnitude of $\frac{d\lambda_+}{dc_{TB}}|_{c_{TB}=0}$ allows us to estimate how different our data will appear for the case with and without a small differentiation. The largest magnitude negative slope we can achieve is -1, when $\beta_B/\delta_T$ approaches zero, so smaller values of $\beta_B$ lead to a larger distinction between the wild and mutant types. Obviously, a larger value of $c_{TB}$ for the wild type will also cause a greater distinction between the wild and mutant infections.

So far, we have assumed that the immune mechanisms are unimportant in determining the values of $I_T(t)$ and $I_B(t)$. However, that does not mean that $Z(t)$ is constant. In fact, we can now plug in our simple analytic solutions for $I_T(t)$ and $I_B(t)$ into (12) to exactly solve for $Z(t)$ at small times. For simplicity, we will only show the solution for the mutant.

$$\frac{dZ^M}{dt} = \left[a_T \frac{I_T(0)e^{\delta t}}{S_0} - \mu\right] Z^M \implies Z^M(t) = Z_0 \exp\left(\frac{a_T}{\delta_T}\frac{I_T(0)}{S_0}(e^{\delta t} - 1) - \mu t\right) \quad (21)$$

This gives us a good idea of how the immune response will be triggered. $I_T(0)/S_0$ is presumably a very small number, so let's assume $a_T I_T(0)/S_0 < \mu$. Then, initially, the number of immune cells decays at a rate $\mu - a_T I_T(0)/S_0$. But as the parasites replicate exponentially, the $a_T \frac{I_T(0)}{S_0} e^{\delta t}$ term will eventually dominate causing rapid growth in $I_T$. The time at which the transition from immune cell decay to immune cell growth occurs is

$$t = \frac{1}{\delta} \log\left( \frac{\mu}{a_T I_T(0)/S_0} \right). \qquad (22)$$

This can be interpreted as the characteristic time at which the immune response is triggered.

### 3. Approach to steady state

The previous section established that whenever (16) is satisfied, infected cell number is driven away from the disease-free equilibrium ($I_T$,$I_B$,$Z$) = (0,0,0). Initially, infected cell number increases exponentially, but immune pressure causes the infected cell number to eventually fall. In this section, we will study the endemic equilibria of the model in order to explain the infection dynamics at late times.

By equating (10), (11), and (12) to zero, three solutions can be found. One is the disease-free equilibrium studied in the previous section, one solution gives a negative value of $Z^*$ and is therefore unphysical, and the final solution is the endemic equilibrium.

$$I_T^* = \frac{\mu}{a_T} \left( 1 - \frac{\frac{a_B}{a_T} c_{TB}}{d_B + \frac{\psi_B}{\psi_T}(\delta_T - c_{TB})} \right) S_0 \qquad (23)$$

$$I_B^* = \frac{\mu}{a_T} \left( \frac{c_{TB}}{d_B + \frac{\psi_B}{\psi_T}(\delta_T - c_{TB})} \right) S_0 \qquad (24)$$

$$Z^* = \frac{\delta_T - c_{TB}}{\psi_T} \left( 1 + \frac{1}{(\delta_T - c_{TB})} \frac{\beta_B c_{TB}}{d_B + \frac{\psi_B}{\psi_T}(\delta_T - c_{TB})} \right) S_0 \qquad (25)$$

The above quantity and all equations in this section are only correct up to linear order in $\frac{c_{TB}}{\delta_T - c_{TB}}$ since the general case is too messy to write down. Of course, if $d_B = \psi_B = 0$ and $c_{TB} > 0$, then there is no way to remove cysts from the system, so $I_B^*$ is infinite, and our solutions are unphysical. So, assume that either $d_B$ or $\psi_B$ is nonzero so that there is some way to remove bradyzoites. It is also useful to consider the ratio of bradyzoite- to tachyzoite-infected cells after a long time

$$\frac{I_B^*}{I_T^*} = \frac{c_{TB}}{d_B + \frac{\psi_B}{\psi_T}(\delta_T - c_{TB})}. \tag{26}$$

In order to study the stability of the endemic equilibrium, we will need to find the eigenvalues and eigenvectors of the Jacobian evaluated at ($I_T$*,$I_B$* ,$Z$*). The Jacobian can be decomposed into the form $J = J_0 + \frac{c_{TB}}{\delta_T - c_{TB}} J_1$, and first-order perturbation theory can be used to find the approximate eigenvalues of $J$. There is a small eigenvalue $\lambda_0 \approx -d_B - \frac{\psi_B}{\psi_T}(\delta_T - c_{TB})$ describing the slow approach to equilibrium for a cyst-dominated system. The inverse of this quantity gives the approximate time it takes for $I_B$ to reach a constant value. The two remaining eigenvalues describe the feedback between infection and immune response.

$$\begin{aligned}\lambda_\pm = &- \frac{c_{TB}(\delta_T - c_{TB})\beta_B}{(d_B + (\delta_T - c_{TB})\frac{\psi_B}{\psi_T})((\delta_T - c_{TB}) + \mu)} \\ &\pm i\sqrt{\mu(\delta_T - c_{TB})}\left[1 + \frac{c_{TB}(\beta_B \mu - \frac{a_B}{a_T}(\delta_T - c_{TB})^2)}{(\delta_T - c_{TB})(d_B + (\delta_T - c_{TB})\frac{\psi_B}{\psi_T})((\delta_T - c_{TB}) + \mu)}\right]\end{aligned} \tag{27}$$

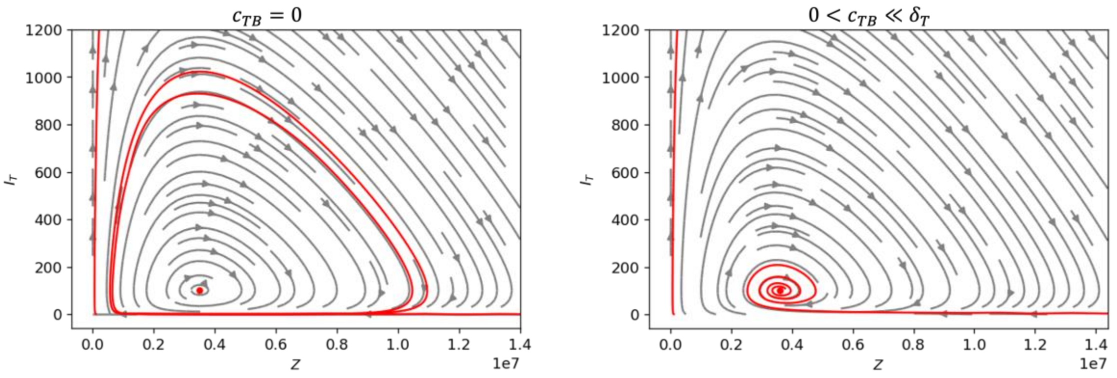


Figure 1: Two example trajectories in $I_T$-$Z$ phase space predicted by equations (10),

(11), and (12) fixed in the plane $I_B = I_B^*$ . Streamlines are shown in gray, and an example trajectory is shown in red leaving the disease-free equilibrium and approaching the steady state $(I_T, I_B, Z) = (I_T^*, I_B^*, Z^*)$ as given by equations (23), (24), and (25).

In the limit $c_{TB} \to 0$, the eigenvalues become purely imaginary $\lambda_\pm = \pm i\sqrt{\mu\delta_T}$. This result suggests that if the mice did not die so quickly, one would expect to see oscillations in infected cell number for the mutant-infected mice around the equilibrium value of $c_{TB} = 0,\ (I_T^*, I_B^*, Z^*) = (\frac{\mu}{a_T}, 0, \frac{\delta_T}{\psi_T})S_0$. This is not surprising because equations (10) and (12) reduce to predator-prey dynamics in that limit. As $I_T$ increases, $Z$ increases which causes $I_T$ to decrease which causes $Z$ to decrease and the cycle continues. Near the critical point, this cycle has a period of

$$T = \frac{2\pi}{|\operatorname{Im}\lambda_\pm|} = \frac{2\pi}{\sqrt{\mu\delta_T}}. \tag{28}$$

Once $c_{TB} > 0$, the period is modified by the bracketed term, but more importantly, the eigenvalues now possess a real part that is always negative for $\beta_B > 0$, indicating a stable focus. So, allowing for conversion between the slow- and fast-replicating forms causes the system to approach the endemic equilibrium with constant infected cell number. Furthermore, as the conversion rates $c_{TB}$ and $\beta_B$ increase, the system reaches equilibrium faster. Using the fact that $(\delta_T - c_{TB}) \gg \mu$, the equilibration time is

$$\tau = -\frac{1}{\operatorname{Re}\lambda_\pm} \approx \frac{d_B + (\delta_T - c_{TB})\frac{\psi_B}{\psi_T}}{c_{TB}\beta_B}. \tag{29}$$

The fact that the having two phases stabilizes the endemic equilibrium is an important observation that we should elaborate on. The simple feedback cycle between $I_T$ and $Z$ is complicated by the fact that some of the tachyzoites are converted into bradyzoites, where they are protected by the immune response. This temporary removal of parasites from the fast-replicating phase dampens the oscillations in $I_T$.

Representative trajectories in phase space are shown in Figure 1. On the left, $c_{TB} = 0$, and solutions leave the disease-free equilibrium and spiral around the center (red dot). On the right, a small $c_{TB}$ value is used, and solutions leave the disease-free equilibrium and spiral toward the stable focus.

Our analysis in this section was assuming a small value of $c_{TB}$ which resulted in under-damped oscillations in phase space. As $c_{TB}$ gets larger, our analysis fails, but the intuition we gained suggests that the solution should approach equilibrium even faster. In section 5, we will see that it is also possible to have over-damped oscillations toward equilibrium when $c_{TB}$ is large enough, indicating purely real negative eigenvalues.

### 4. Numerical values

Given a combination of experimental data and the intuition we've obtained from this analysis, we can come up with a reasonable range of parameters to explore. Formulas will be given to show how each parameter's mean and error were calculated. All rates are given in units of $d^{-1}$.

The dynamics are greatly simplified for the mutant where the initial growth rate is governed by just a single parameter $\delta_T$. Assuming immune clearance is negligible for the first two data points, we can extract $\delta_T$ from the infection curve assuming the form (18).

$$\delta_T = \frac{1}{t_2 - t_1}\left[\log\left(\frac{I_T^M(t_2)}{I_T^M(t_1)}\right) \pm \sqrt{\left(\frac{\sigma_{I_T^M(t_1)}}{I_T^M(t_1)}\right)^2 + \left(\frac{\sigma_{I_T^M(t_2)}}{I_T^M(t_2)}\right)^2}\right] \tag{30}$$

Using the full data set (compiling all of the data for both trials of the experiment) gives $\delta_T = .711 \pm .238$. Note that the rates $\beta_T$ and $d_T$ only appear as a difference in the equations for $I_T$ and $I_B$, so we cannot extract these quantities individually from the infection curves. However, we know that the doubling rate of tachyzoites once in a cell is about six hours, and a single parasite doubles about four times before bursting. Thus, the mean lifetime of a $T$-infected cell is about $d_T^{-1} = 1$ day and $n_T = 16$. To get an idea for the error, we can consider the possibility that the parasites double three or five times, leading to a range $d_T^{-1} \in [.75, 1.25]$ and $n_T \in [8,32]$. Using our extracted value of $\delta_T$ and the known values of $d_T$ and $n_T$, we can estimate $\beta_T \approx 1.7$ and $f_{T0} \approx .1$.

Next, we can apply the inequality (20) to place a bound on $\beta_B - d_B$. First note that $\beta_B - d_B = (n_B f_{B0} - 1)d_B \gg d_B$. In words, when a cyst bursts, the number of parasites which go on to infect new cells is much greater than one. Thus, we have $d_B \ll \beta_B < \delta_T$. Although it is well-known that the dynamics of bradyzoites is much slower than that of tachyzoites (we estimate $d_B^{-1}$ >50 days), the magnitude of $\beta_B = d_B n_B f_{B0}$ is less clear since $n_B$ can be in the hundreds or thousands. Using our value for $\delta_T$, we find that the time scale over which bradyzoites generate new tachyzoites is $\beta_B < 1$. Although this doesn't forbid the possibility of $\beta_B$ being close to 1, the fact that we see such a large difference between the wild and mutant infection curves (nearly four times as many tachyzoite-infected cells in the mutant by day 5) suggests that $\beta_B$ is likely very small, at least early on. Setting both $d_B$ and $\beta_B$ equal to zero in (17) for simplicity allows us to extract $c_{TB}$ from our data.

$$c_{TB} = \frac{1}{t_2 - t_1}\left[\log\left(\frac{I_T^M(t_2)}{I_T^M(t_1)}\frac{I_T^W(t_1)}{I_T^W(t_2)}\right) \pm \sqrt{\left(\frac{\sigma_{I_T^M(t_1)}}{I_T^M(t_1)}\right)^2 + \left(\frac{\sigma_{I_T^M(t_2)}}{I_T^M(t_2)}\right)^2 + \left(\frac{\sigma_{I_T^W(t_1)}}{I_T^W(t_1)}\right)^2 + \left(\frac{\sigma_{I_T^W(t_2)}}{I_T^W(t_2)}\right)^2}\,\right] \quad (31)$$

Using the full data set gives a value of $c_{TB} = .227 \pm .360$. It may seem strange that negative values of $c_{TB}$ appear here which are physically impossible, but this arises from how we have calculated $c_{TB}$. We are assuming that the parasites replicate exponentially between times $t_1$ and $t_2$ and that the change in the separation between the wild and mutant infection curves can be explained by the parameter $c_{TB}$. But what we saw in the first data set is that for some data points, the ratio $I_T^M(t_1)/I_T^W(t_1)$ is bigger than the ratio $I_T^M(t_2)/I_T^W(t_2)$, indicating that the growth rate of the parasites in the wild type is actually greater than that of the mutant. This could indicate that by day 5, the immune response is already large enough that our linear model fails, but without more data at early times, we can't say for sure. Allowing for a nonzero value of $\beta_B < \delta_T$ does not cause the predicted negative $c_{TB}$ values to become positive, but it does increase the magnitude of the predicted $c_{TB}$ values by a small amount that is still within our predicted error bounds. Enforcing a positive value for $c_{TB}$, we find that $c_{TB} \in [0,.587]$, but most likely, it's closer to .2.

Estimating the values for the immune response is much more challenging. We must stress that this mathematical model should not be thought of as a fit to data. Once the immune mechanisms are comparable to the replication/conversion mechanisms, the system is too complex to study quantitatively with so few data points. Here, we just want to determine a reasonable range of parameters to study numerically. First, let's estimate the values of $a_T$, $\psi_T$, and $\mu$ which govern the immune response of the mutant. We already assumed that by $t = 5$, the $\psi_T \frac{I_T Z}{S_0}$ term in equation (10) is negligible, which allowed us to use the approximate solution (18). Similarly, by $t = 5$, we should be able to use (21) to estimate the growth of immune cells.

Except for when $I_T(t)$ is very small, the $\mu t$ term is negligible, and we have

$$\frac{a_T}{S_0} \approx \frac{\delta_T}{I_T(t_2) - I_T(t_1)} \log\left(\frac{Z(t_2)}{Z(t_1)}\right) \left[1 \pm \sqrt{\left(\frac{\sigma_{\delta_T}}{\delta_T}\right)^2 + \frac{\sigma^2_{I_T(t_1)} + \sigma^2_{I_T(t_2)}}{(I_T(t_2) - I_T(t_1))^2} + \left(\frac{1}{\log(Z(t_2)/Z(t_1))}\right)^2 \left(\frac{\sigma^2_{Z(t_1)}}{Z(t_1)^2} + \frac{\sigma^2_{Z(t_2)}}{Z(t_2)^2}\right)}\right] \quad (32)$$

Plugging in our results, we find $a_T/S_0 = .00101 \pm .00098$. The parameter $\mu$ is important in dictating the relative magnitude of $I_T$ and $Z$. Notice by equating (12) to zero that when $Z(t)$ is at a maximum or

| Symbol | Description | Numerical estimate |
|---|---|---|
| $d_T$ | Death rate of *T* cells (without any immune response) | .8 - 1.3 $d^{-1}$ |
| $\delta T$ | Growth rate of *T* cells (prior to immune response) | .5 - 1 $d^{-1}$ |
| $c_{TB}$ | Tachyzoite to bradyzoite conversion rate | 0 - .6 $d^{-1}$ |
| $\beta B$ | Bradyzoite to tachyzoite re-activation rate | $< \delta_T$ |
| $dB$ | Death rate of *B* cells (without any immune response) | $\ll \beta_B$ |
| $aT/S0$ | Tachyzoite-induced immune activation rate | 0 - .002 $d^{-1}$ |
| $aB/S0$ | Bradyzoite-induced immune activation rate | $\ll a_T/S_0$ |
| $\mu$ | Immune cell death rate | $(1 - 100) * a_T/S_0$ |
| $\psi T/S0$ | Tachyzoite immune killing rate | $(0.8 - 2.4) * 10^{-7}$ $d^{-1}$ |
| $\psi B/S0$ | Bradyzoite immune killing rate | $\ll \psi_T/S_0$ |

Table 1: Parameters used in the paper. One final parameter, the contact rate $\beta_T = \delta_T + d_T$ is sometimes used instead of $\delta_T$.

minimum, $I_T$ satisfies $I_T(Z = Z_{\max}) = \frac{\mu}{a_T/S_0}$. However, using our full data set, $Z(t)$ is a monotonically increasing function, indicating that $I_T(Z = Z_{\max})$ is smaller than any $I_T$ reached in this experiment. We don't know how small $I_T$ needs to be before the immune cells start disappearing, but we can at least say $I_T(Z = Z_{\max}) < 100$. But, one would expect that if only a single parasite was introduced, the immune response would not be triggered until the parasite begins replicating. This allows us to bound $\mu$.

$$\frac{a_T}{S_0} < \mu < 100\frac{a_T}{S_0} \tag{33}$$

Or, using our mean value for $a_T/S_0$, we find $\mu < .1$. Lastly, we can determine $\psi_T$ using a similar trick to how we estimated $\mu$. By setting (10) to zero, we find that the value of $Z$ where $I_T$ takes its maximum value must satisfy $Z(I_T = I_{T,\max}) = \frac{\delta_T}{\psi_T/S_0}$. Unlike when we estimated $\mu$, we actually see the peak in $I_T$ in our experiment around $t = 11$ days, so we can explicitly use the value $Z(I_T = I_{T,\max}) = Z(t = 11)$. Thus, we have

$$\frac{\psi_T}{S_0} = \frac{\delta_T}{Z(I_T = I_{T,\max})}\left[1 \pm \sqrt{\left(\frac{\sigma_{\delta_T}}{\delta_T}\right)^2 + \left(\frac{\sigma_{Z(I_{T,\max})}}{Z(I_T = I_{T,\max})}\right)^2}\right]. \tag{34}$$

Plugging in these numbers gives $\psi_T/S_0 = 1.62 * 10^{-7} \pm 0.79 * 10^{-7}$.

Lastly, we have the parameters $a_B$ and $\psi_B$ governing the immune response to bradyzoites. While little is known about this immune mechanism, it is safe to say that $a_B \ll a_T$ and $\psi_B \ll \psi_T$ since it is harder to detect and defeat the encysted cells.

A summary of the parameters in the model is given in table 1. Additionally, we have four initial conditions which we will estimate as $S_0 = 10^8$, $I_T(0) = 1$, $I_B(0) = 0$, and $Z(0) = 10^5$.

### 5. Simulations

#### a. No immune response

An important observation in the data is that the separation between the wild-type and mutant variants happens within days of reaching the brain. In order to explain this immediate difference, we consider first the linear model without immune response which is a good approximation of the full solution at early times. The parameter $\delta_T$ simply controls the growth rate. Although this is perhaps the single most important parameter in setting the scale of the infection, mathematically it is the easiest parameter to understand. The parameter $d_B$ is so small that setting it equal to zero does not change the dynamics significantly at early times, so we fix $d_B = 0$. That leaves two interesting rates to investigate: $c_{TB}$ and $\beta_B$. Numerical results are shown in figure 2. The mean values of $\delta_T$ and $d_T$ from the previous section are used. Three values of $c_{TB}$ are used: $c_{TB} = 0$ corresponds to the mutant, $c_{TB} = .25$ is a reasonable estimate for the wild type, and $c_{TB} = .75$ is used to investigate the hypothetical *T. gondii* variant that has an abnormally fast conversion rate. When $\beta_B < \delta_T$, the infection is more severe for the mutant, as is seen in the experiment. When $\beta_B$ is close to $\delta_T$, the value of $c_{TB}$ is unimportant. When $\beta_B > \delta_T$, the infection is more severe for the wild type. All of these numerical results agree with our analytic predictions in section 2.

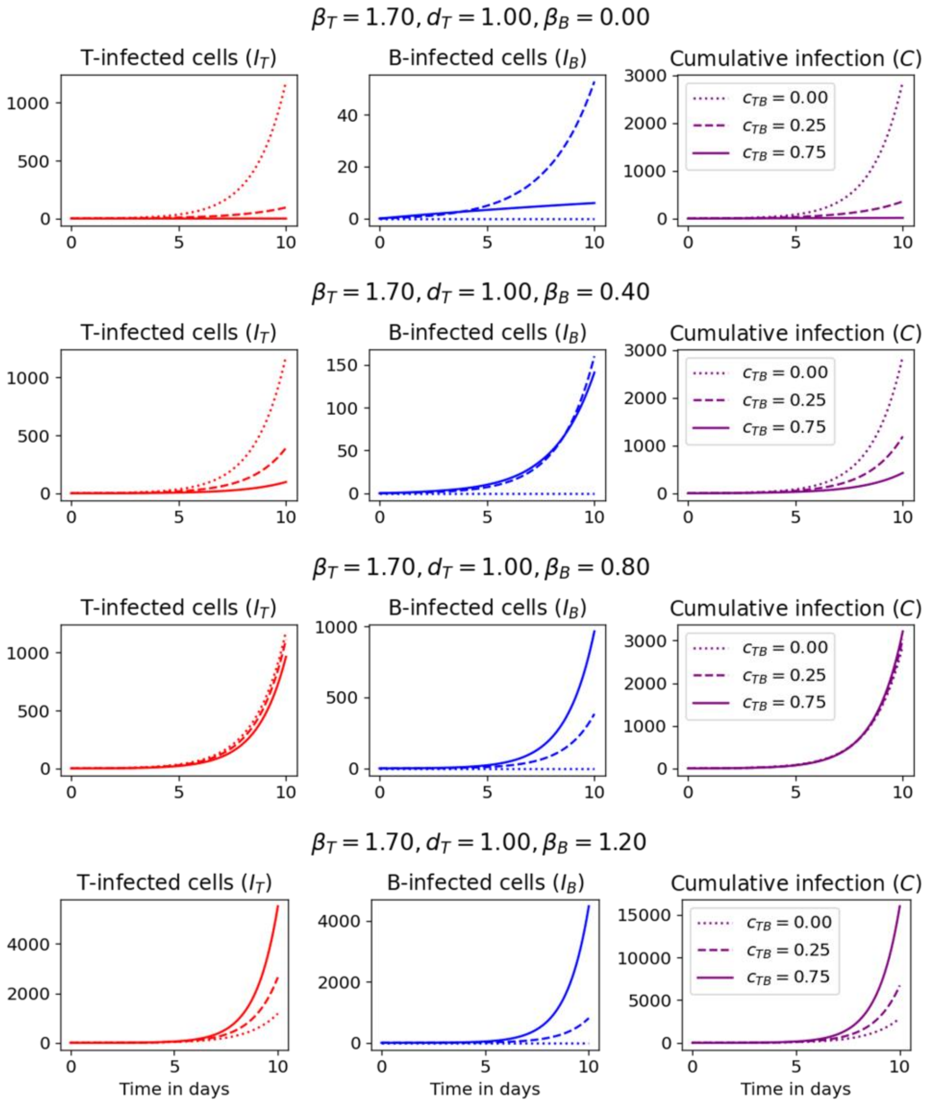


Figure 2: Results to equations (10), (11), and (13) with all immune-response parameters set to zero.

#### b. With immune response

Because we have eliminated the nonlinearity arising from a finite number of cells, the only feedback mechanism in our problem is the immune response. So, in order to limit the number of infected cells and obtain reasonable results after about 5 days of infection, it is important that we include the parameters governing the immune response. Numerical results are given in figure 3.

In all cases, there is an initial exponential growth of infections which is suppressed by the immune response. In the first four plots, the bradyzoite immune response is turned off. When $c_{TB}$ = 0 (top row), the immune cells can remove all of the infected cells. Once most tachyzoite-infected cells are removed, the immune cells die at a rate of $\mu$, but this decrease in immune cells promotes further tachyzoite spread. This feedback loop causes periodic solutions in $I_T$. When a small value of $c_{TB}$ is introduced (second row), oscillations are observed at a period given by (28), but the strength of the infections decays with time. Both of these curves are consistent with the analysis given in section 3. In the third and fourth rows, $c_{TB}$ is increased to the point where our analysis from section 3 fails. The endemic equilibrium becomes a stable node. The infection dynamics are overdamped, and depending on the size of $d_B$, $I_B$ may either increase monotonically or reach a peak and then decrease to its steady state value. In the fifth row, a small bradyzoite immune response is introduced and underdamped solutions are again observed. Increasing the strength of the bradyzoite immune response (sixth row) increases the relaxation time, and large oscillations are observed for a longer period of time. This is at least qualitatively in agreement with equation (29) which suggests that a larger bradyzoite immune response should increase the relaxation time.

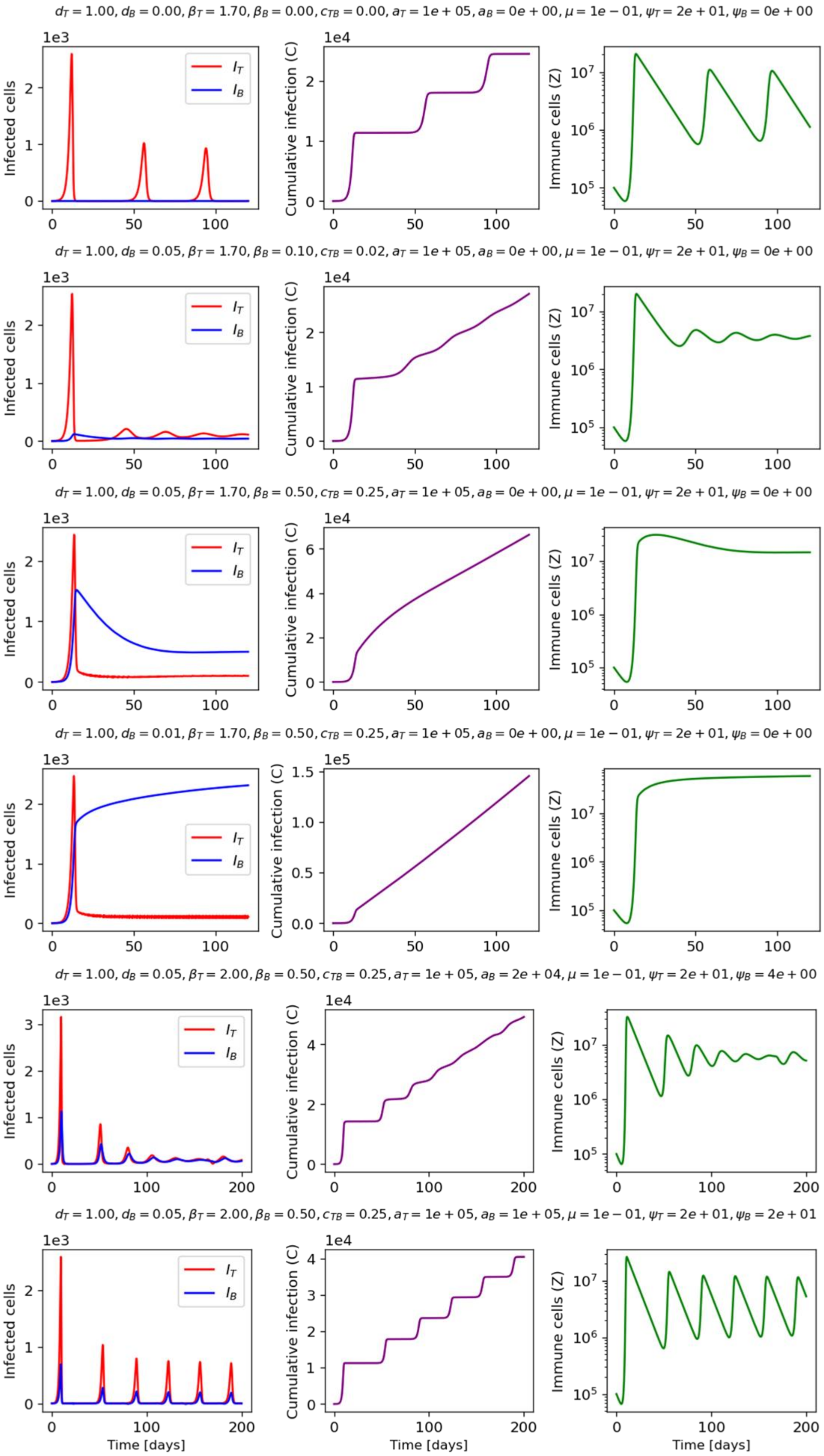

$d_T = 1.00, d_B = 0.00, \beta_T = 1.70, \beta_B = 0.00, c_{TB} = 0.00, a_T = 1e+05, a_B = 0e+00, \mu = 1e-01, \psi_T = 2e+01, \psi_B = 0e+00$
$d_T = 1.00, d_B = 0.05, \beta_T = 1.70, \beta_B = 0.10, c_{TB} = 0.02, a_T = 1e+05, a_B = 0e+00, \mu = 1e-01, \psi_T = 2e+01, \psi_B = 0e+00$
$d_T = 1.00, d_B = 0.05, \beta_T = 1.70, \beta_B = 0.50, c_{TB} = 0.25, a_T = 1e+05, a_B = 0e+00, \mu = 1e-01, \psi_T = 2e+01, \psi_B = 0e+00$
$d_T = 1.00, d_B = 0.01, \beta_T = 1.70, \beta_B = 0.50, c_{TB} = 0.25, a_T = 1e+05, a_B = 0e+00, \mu = 1e-01, \psi_T = 2e+01, \psi_B = 0e+00$
$d_T = 1.00, d_B = 0.05, \beta_T = 2.00, \beta_B = 0.50, c_{TB} = 0.25, a_T = 1e+05, a_B = 2e+04, \mu = 1e-01, \psi_T = 2e+01, \psi_B = 4e+00$
$d_T = 1.00, d_B = 0.05, \beta_T = 2.00, \beta_B = 0.50, c_{TB} = 0.25, a_T = 1e+05, a_B = 1e+05, \mu = 1e-01, \psi_T = 2e+01, \psi_B = 2e+01$
Infected cells
Cumulative infection (C)
Immune cells (Z)
$I_T$
$I_B$
Time [days]

### **References**


[1] Adam Sullivan, Folashade B. Agusto, Sharon Bewick, Chunlei Su, Suzanne Lenhart, and Xiaopeng Zhao. A mathematical model for within-host toxoplasma gondii invasion dynamics. *Mathematical biosciences and engineering : MBE*, 9:647–62, 07 2012.

[2] Mark H. Holmes. *Introduction to perturbation methods*. Texts in applied mathematics. Springer, New York, 2nd ed edition, 2013.
